# MetaSAG: A Tool for Multi-level Exploration and Taxonomic Analysis of Microbial Single-Amplified Genomes

**DOI:** 10.64898/2026.03.04.709444

**Authors:** Sainan Zhang, Meiyu Du, Guanzhi He, Kai Qian, Kexin Li, Baifeng Li, Ping Wang, Minke Lu, Xiaoliang Wu, Chao Wang, Hongbin Han, Shuofeng Yuan, Xue Zhang, Liang Cheng

## Abstract

Microbial single-amplified genome (SAG) sequencing technologies have elevated microbial research resolution to the single-cell level. However, neither upstream data processing nor downstream analysis has been fully developed, greatly limiting the research in strain level. Herein, we developed **MetaSAG** (**Multi-level Exploration and Taxonomic Analysis of microbial Single-Amplified Genomes**), which enables accurate and rapid taxonomic classification of microbial SAGs. MetaSAG outperforms existing method in species classification certainty, computational efficiency, and sensitivity of low abundance species identification. In addition, MetaSAG enables species-level functional analysis, as well as strain-level evolutionary analysis. With the help of MetaSAG, we discovered the parasitic relationship between phages and bacteria, identifying multiple susceptible bacteria and a broad spectrum of phages. Furthermore, we developed **MetaK-Lytic** (**k-mer-based meta-learning framework to predict phage lytic ability**) to achieve accurate prediction of phage lytic activity based on 31-mer short sequences, which is well adapted to the characteristics of incomplete SAG sequences. Overall, we offer a comprehensive integrated tool that can parse microbial SAG data from raw data to the strain level to decipher the functional ecology of microbial dark matter, with broad implications for microbial ecology and phage therapy (https://github.com/liangcheng-hrbmu/MetaSAG).

**Significance Statement:** This work provides a comprehensive framework for high-resolution SAG data analysis. The developed pipeline improves taxonomic annotation sensitivity and speed. Strain-level tracking enables dynamic evolutionary and functional insights, while single-cell bacterial-virus network reconstruction reveals precise interaction patterns. The novel annotation-free short sequence-based MetaK-Lytic facilitates functional prediction of uncharacterized phage sequences. Integrated into the MetaSAG platform, these tools deliver a streamlined, multi-level solution for interpreting SAG data, advancing studies in microbial ecology, evolution, and virology.

## Introduction

The human intestinal tract harbors a complex microbial ecosystem that plays a pivotal role in maintaining host health, including regulating immune function ^1^, promoting nutrient absorption ^2^, and preserving intestinal barrier integrity ^3^. To characterize these communities, conventional sequencing approaches such as 16S rRNA gene sequencing and metagenomic sequencing have been widely adopted. However, these methods possess inherent limitations. 16S sequencing often fails to resolve taxonomic units beyond the genus level and is susceptible to amplification biases ^4^, while metagenomic sequencing, which relies on bulk DNA extraction, can be dominated by abundant species, thereby obscuring signals from low-abundance taxa ^5^. Furthermore, both techniques generally lack the resolution to distinguish strain-level heterogeneity—a critical dimension for understanding functional diversity, pathogen virulence ^6,7^, and antibiotic resistance ^8^. And most importantly, they cannot directly link genetic elements to their cellular hosts, impeding research into microbe-host interactions.

The recent advent of microbial single-amplified genome (SAG) sequencing, particularly Microbe-seq ^9^, represents a transformative advance. By isolating and sequencing thousands of individual microorganisms within microfluidic droplets, this technology provides direct access to individual genomes, bypassing the biases introduced by bulk DNA extraction. This approach significantly improves the detection of rare species and “microbial dark matter”, avoids assembly chimerism, and most importantly, enables the direct analysis of genomic variation at the strain level.

Despite its transformative potential, the analysis of Microbe-seq data remains challenging. The low genomic coverage typical of single-cell data has necessitated reliance on complex, multi-step clustering and co-assembly protocols based on genomic similarity ^9^. These processes are computationally intensive, exhibit poor reproducibility, especially with large datasets, and struggle to robustly identify technical artifacts like multi-species droplets or low-abundance species. An alternative strategy is the taxonomic classification of droplets based on short-read annotations, which places stringent demands on the reference database: it requires comprehensive coverage, high specificity of marker genes, annotation accuracy, and computational efficiency. Existing solutions each have shortcomings; for instance, the 16S databases used by QIIME2 ^10^ offer limited coverage, while whole-genome alignment tools like Kraken2 ^11^ and Kaiju ^12^ are prone to false assignments due to sequence homology. In contrast, MetaPhlAn4 ^13^ achieves superior specificity by utilizing a curated set of clade-specific marker genes from over 20,000 species, offering a promising path for accurate single-cell taxonomic profiling.

Beyond bacteria, the gut virome, dominated by bacteriophages (phages), is a key regulator of microbial ecosystems. Phages are the most abundant biological entities in the human gut. Cohort studies have demonstrated that the gut phage community is highly dynamic ^14^, and phage predation is a powerful driver of bacterial population diversity. The lytic capability of phages is already being harnessed in clinical settings for treating drug-resistant bacterial infections ^15–17^, underscoring their therapeutic potential. However, a comprehensive understanding of in situ phage-host interaction networks, including host range and life cycle states (lytic vs. lysogenic), has been hindered by the limitations of bulk sequencing methods.

Besides, current computational methods for phage lytic ability prediction rely heavily on the presence of known marker genes ^18,19^ (e.g., integrases, immunity repressors), which fails for phages lacking these characterized genes or for novel phage families. This limitation is exacerbated in the context of SAG sequencing, which primarily captures uncultured and previously unknown phages, creating a scenario of extreme data scarcity where traditional supervised learning models are prone to overfitting.

Inspired by these challenges, we are committed to developing a comprehensive and integrated upstream and downstream analysis process for microbial SAG data, including analysis at the species and strain levels, as well as the interaction between bacteriophages and hosts, to fully understand the community structure, function, dynamic evolution of intestinal microorganisms, the prediction of bacteriophage lysis ability and so on.

## Results

### 1. Development and performance benchmarking of SAG analysis pipeline

#### 1.1 Data Quality Control

High-throughput single-cell amplified genome sequencing (Microbe-seq) generates a substantial fraction of droplets with limited reads, complicating robust taxonomic annotation. In the methodology established by Zheng et al. ^9^, after ordering the droplets according to the number of reads contained in them, the first 5% and the last 15% of the droplets were removed. While this uniform filtering strategy can remove some multicellular or low-quality droplets, it is suboptimal because it applies a fixed percentile cutoff regardless of the underlying read count distribution, which can vary substantially between runs. To overcome this limitation, we established a statistically-grounded and objective quality control threshold (r*) by identifying the inflection point (i.e., the point of maximum curvature) in the cumulative distribution of droplet read counts (**Fig. 2A, B**, see **Materials and methods**). This method locates a natural breakpoint in the data that separates low-coverage droplets from the main population in a reproducible and dataset-specific manner.

**Figure 1.**
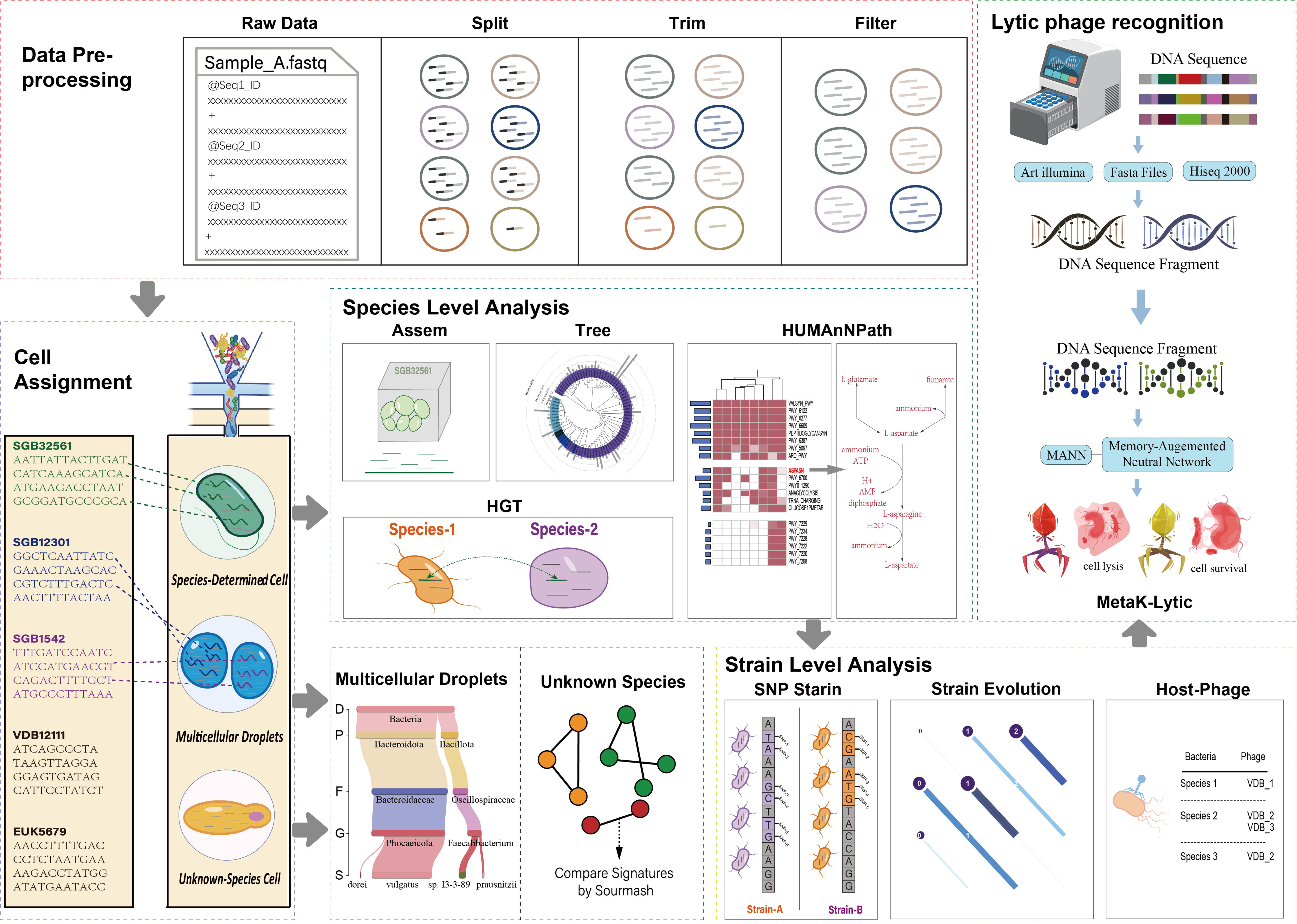
**The overview and main modules of MetaSAG.**

**Figure 2.**
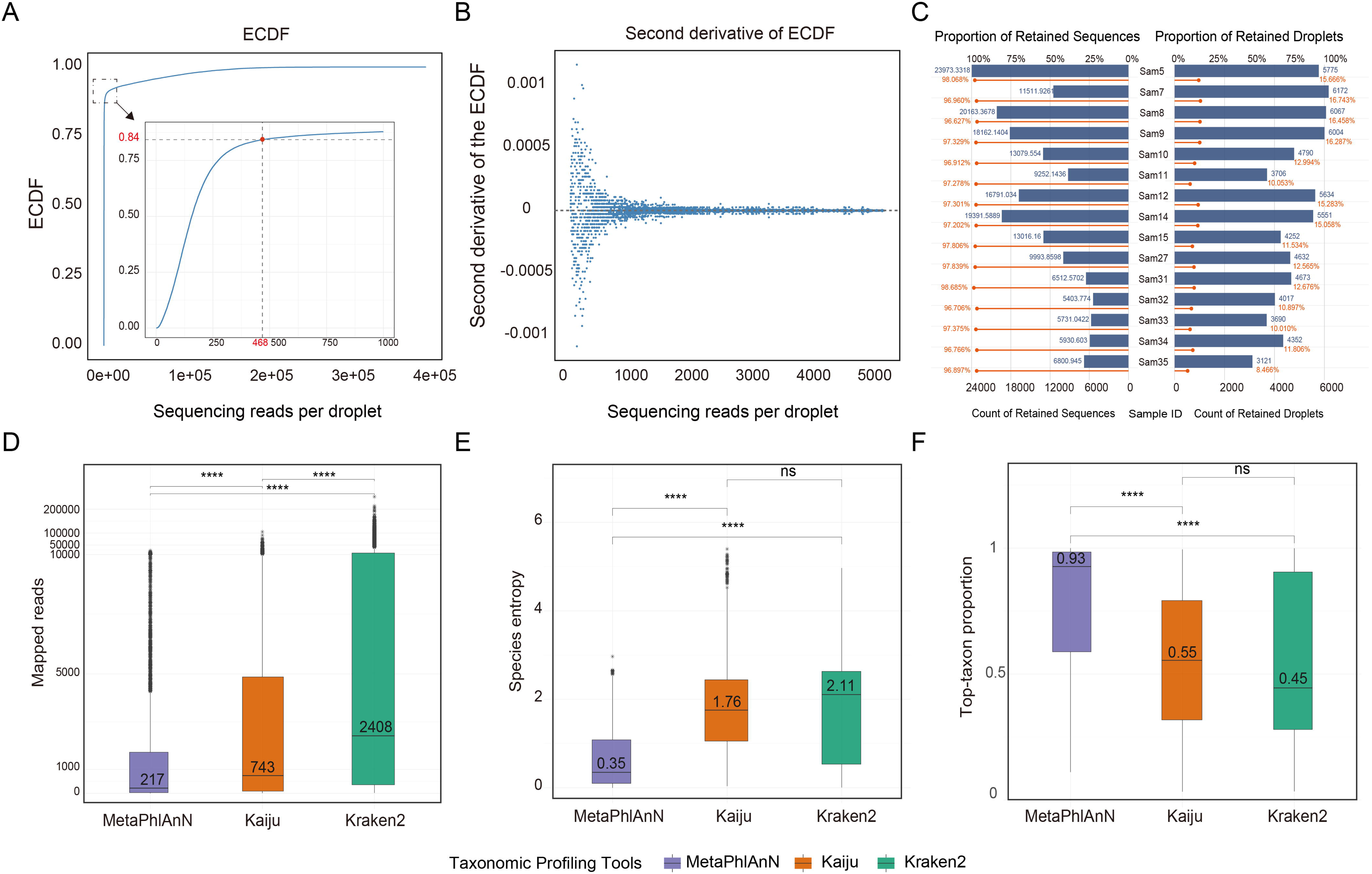
Performance of MetaSAG in species annotation. **(A)** Cumulative distribution curve of read counts per droplet in Sample 5. **(B)** Second derivative of the cumulative distribution of droplet read counts in Sample 5. **(C)** Number (blue bars) and proportion (red line) of retained reads and droplets per sample. **(D)** Number of mapped reads obtained by each alignment tool. **(E)** Entropy of species assignments per droplet calculated for each of the three alignment tools. **(F)** Proportion of reads assigned to the dominant taxon by each alignment tool.

Application of this data-driven threshold filtered out 83.25–91.53% of droplets, which were predominantly empty or contained minimal genetic material. Crucially, it conserved 96.62–98.68% of the sequencing reads (**Fig. 2C**). This efficient separation of signal from noise provided a solid basis for downstream analyses.

### 1.2 The selection of alignment tools

In order to select a reliable classification tool as the core of SAG pipeline, we next benchmarked the performance of three representative taxonomic classifiers—Kraken2 (k-mer based), Kaiju (translation-based), and MetaPhlAn4 (marker-gene based)—on the filtered dataset. To evaluate classification confidence, we calculated the entropy of species-level assignments per droplet to quantify the degree of classification confusion of each tool in single-droplet annotation. The results showed that MetaPhlAn4 mapped a relatively small total number of reads (**Fig. 2D**) but had significantly lower entropy of droplets than both Kaiju (P = 4.1×10^-5^, **Fig. 2E**) and Kraken2 (P = 2.3×10^-6^, **Fig. 2E**), indicating that MetaPhlAn4 may have superior classification certainty while use the relatively less sequence information. Consistent with this, the median fraction of reads assigned to the dominant taxon was 0.93 for MetaPhlAn4, substantially higher than that for Kraken2 (median = 0.55, P = 1.8×10^-7^) and Kaiju (median = 0.45, P = 3.2×10^-6^) (**Fig. 2F**), confirming its more accurate species discrimination ability.

These performance advantages are attributable to the analytical strategy of MetaPhlAn4, which leverages clade-specific marker genes to minimize cross-mapping among related species and disregards non-informative genomic regions, thereby enhancing specificity and confidence in taxonomic calls. Given the superior performance, we choose MetaPhlAn4 as the core classifier for our pipeline (MetaSAG).

### 1.3 Comparison with existing methods

#### Concordance Validation: MetaSAG achieves high concordance with established taxonomic assignment methods

To assign confident taxonomic identities, we designated the species with the highest annotation proportion in each droplet as its “primary species”. Then MetaSAG was rigorously benchmarked against two methods: (1) genome similarity-based clustering with co-assembly, and (2) metagenomics-based taxonomy. Comparison with genome clustering revealed a discordance rate of only 5.9% (758 out of 12,957 droplets; **Fig. 3A**). When evaluated against metagenomics taxonomy, the discordance was even lower at 5.0% (60 out of 1,198 unambiguously assigned droplets; **Fig. 3A**). We further compared the species identification performance between the method described by Zheng et al. ^9^ and metagenomics. Although the intersection agreement between their method and metagenomics data was slightly higher, it identified fewer total species than MetaPhlAn4-based method (**Supplementary Fig. 1**). Collectively, these benchmarks demonstrate that MetaSAG, as a marker gene-based classification tool, it can accurately recapitulate species-level taxonomy while providing more comprehensive species recovery.

**Figure 3.**
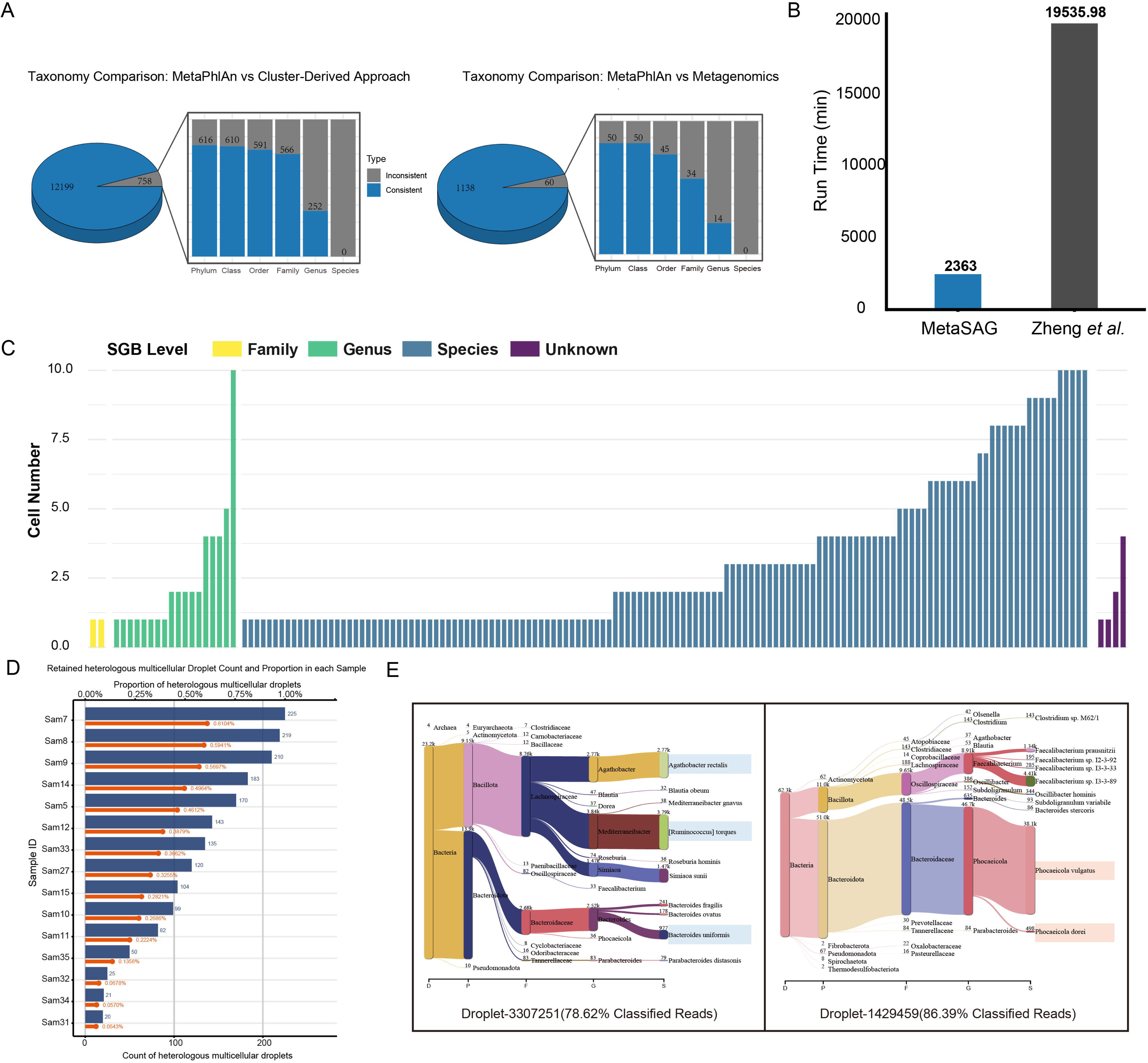
The Advantages of MetaSAG. (**A**) The consistent and inconsistent droplets/species that MetaSAG intersected with Cluster-Derived Approach (Zheng’s method) and Metagenomics. Blue indicates agreement and gray indicates disagreement. The bar graph represents the distribution of agreement at each taxonomic level for species with inconsistent classification. (**B**) The time consumption of MetaSAG and Zheng’s method. (**C**) The distribution of low abundance species identified by MetaSAG and their taxonomic level. (**D**) Distribution of heterologous multicellular droplets after filtration using the threshold (20%). (**E**) Heterologous multicellular droplets annotated by Kracken2. The figure shows the four species contained in the droplet and the relationships among these species.

#### Runtime Performance: MetaSAG achieves significant acceleration over existing co-assembly methods

We directly compared the computational time required by MetaSAG against the established co-assembly and clustering pipeline described by Zheng *et al.* ^9^, using an identical dataset of 15 samples. On a standard computing node (96 CPU cores, 1024 GB RAM), MetaSAG completed species annotation and genome assembly in about 39 hours. In stark contrast, the co-assembly-based method required over 13 days to process the same data (only assembled and without annotations), representing an at least 87.9% reduction in wall-clock time (**Fig. 3B, Supplementary Table 1**). This dramatic acceleration stems from MetaSAG’s streamlined, classification-first architecture, which bypasses the computationally intensive steps of individual genome assembly, cross-sample clustering, and iterative co-assembly.

Consequently, MetaSAG transforms single-cell microbiome analysis from a protracted, resource-intensive endeavor into a rapid, scalable workflow, enabling timely insights from large-scale SAG studies. This efficiency translates to practical utility: analyses that previously required weeks of computation can now be completed within 1 or 2 days, dramatically accelerating the feedback loop between experiment and discovery.

#### Low-Abundance Recovery: MetaSAG enables unprecedented access to microbial “dark matter”

We detected 157 low-abundance taxa (each represented by <10 cells) across 470 droplets: 137 taxa were confidently assigned to the species level, 18 to the genus level, and only two remained at the family level (*Lachnospiraceae* and *Ruminococcaceae*; (**Fig. 3C, Supplementary Table 1**). Remarkably, these low-abundance taxa constituted 54.6% of all single-amplified genome bins (SGBs) identified. Given that each droplet represents an independent genomic sample, this extensive catalog of rare lineages underscores the unique power of single-droplet genomics to capture strain-level diversity often obscured in bulk sequencing. Thus, MetaSAG establishes a new technical paradigm for mapping the rare biosphere, transforming our capacity to explore the diversity and ecological roles of microbial “dark matter” within complex communities.

#### 1.4 Identification of heterologous multicellular droplets

In single-cell whole-genome amplification sequencing, the co-encapsulation of distinct microbial species within the same droplet can lead to chimeric assemblies and confound downstream analysis. To reveal this, we formulated an inspection strategy. For a single droplet with more than 200 marker gene reads, if the proportion of marker gene annotations of more than one species exceeds 20% of the total annotation reads in the droplet, the droplet is determined to be a “possible heterologous multicellular droplet”. Applied this strategy across 15 samples, we find that the proportion of heterologous multicellular droplets was below 1% (**Fig. 3D**), indicating the relatively high single-cell encapsulation efficiency of microfluidic sequencing technology.

Among these heterologous multicellular droplets, there comprised 1,724 two-species, 62 three-species, and 2 four-species droplets. Notably, more than 1/3 of these droplets contain species belonging to the same family, suggesting phylogenetically related microbes may be more prone to physical co-aggregation and subsequent co-encapsulation. We then performed independent taxonomic re-annotation on the two four-species droplets using Kraken2 and MetaPhlAn4 to further investigate this heterogeneity (**Fig. 3E and Table 1**). In Droplet-3307251, species identified by MetaPhlAn4 such as *Mediterraneibacter torques*, *Bacteroides uniformis* and *Agathobacter rectalis* also accounts for a significant proportion in the Kracken2 annotation results. Meanwhile, *Acetatifactor intestinalis* and *Simiaoa sunii* were specifically recognized by MetaPhlAn4 and Kracken2, respectively. The annotation result of Droplet-1429459 is the same.

**Table 1.**
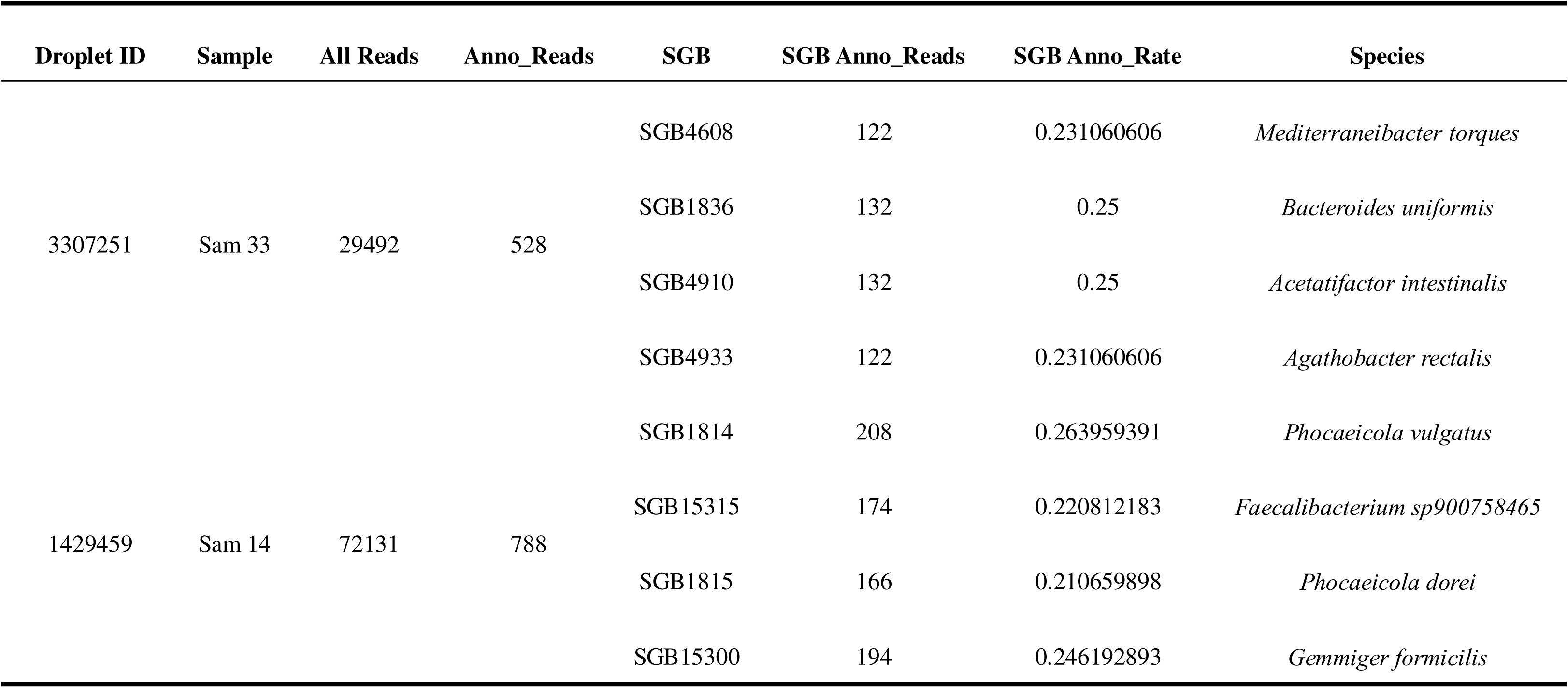
Annotated results of four-species droplets by MetaPhlAn4.

In conclusion, the differences among various annotation tools highlight the inherent distinctions in the reference databases and classifier analysis algorithms they rely on. However, it is clear that this annotation strategy based on MetaPhlAn4 allows us to identify heterologous multicellular droplets and explore their distribution characteristics, providing methodological support and data guarantee for the subsequent analysis of the structure and function of microbial communities.

### 2. Profiling of Microbial Community Structure, Phylogeny, and Function at Single-Cell Resolution

#### 2.1 Ecological insights into community structure

To enhance accuracy in species assignments, we further established a dual-filtering criterion, requiring each droplet to possess >100 total annotated reads and >80% of reads assigned to its primary species (**Supplementary Table 1**). Validation via alignment to reference genomes showed that droplets passing these filters achieved a mean alignment rate exceeding 50%, significantly higher than those do not meet the threshold (**Fig. 4A**), confirming the efficacy of this method in excluding low-confidence annotations.

**Figure 4.**
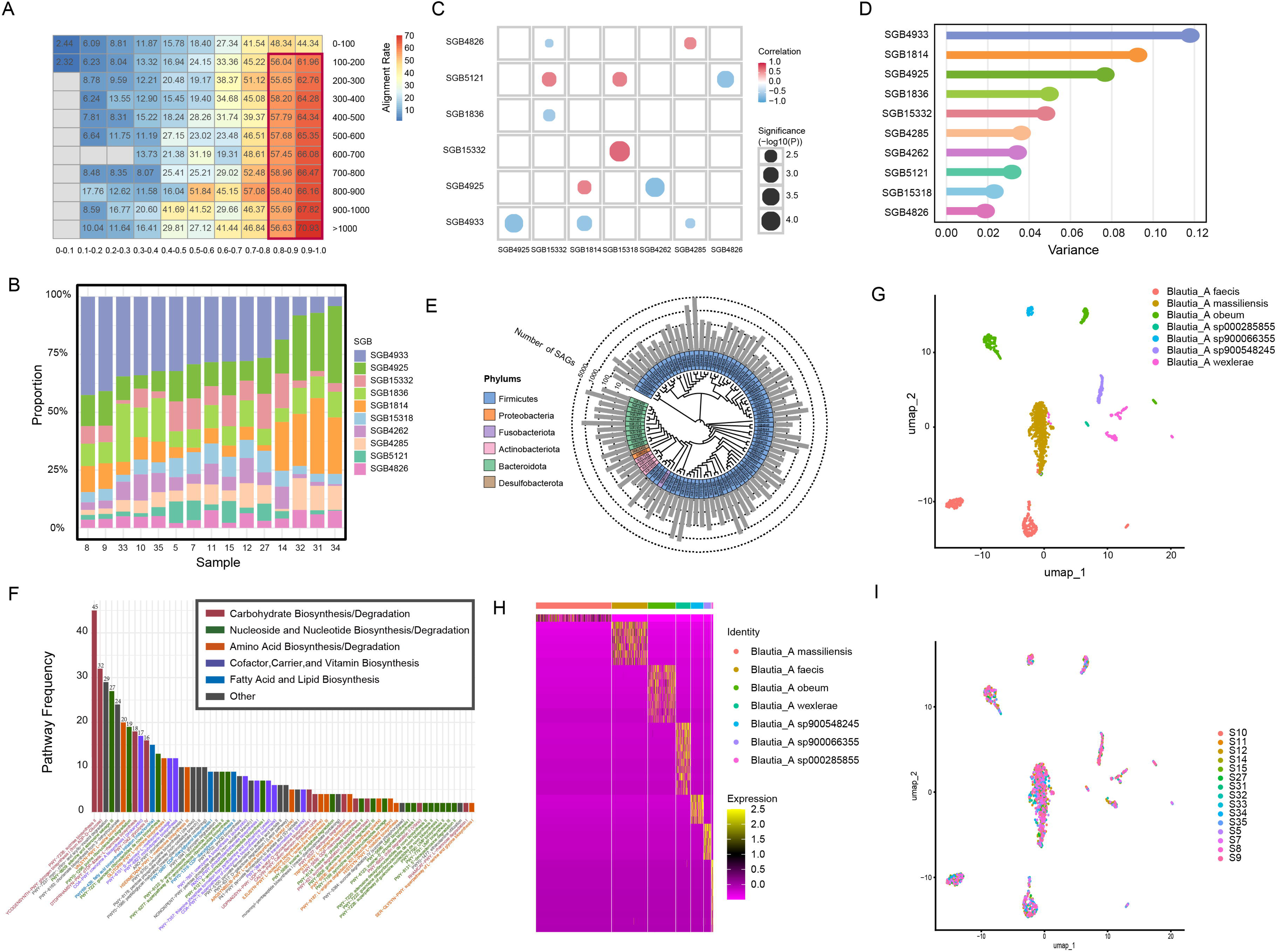
Analysis of community structure in the human gut. (**A**) Average distribution of alignment rates for the dominant species within droplets. Rows represent read counts, columns represent alignment rates, and colors correspond to alignment rate. (**B**) Stacked bar plot showing the relative abundance of top 10 species in each sample. (**C**) Bubble chart illustrating correlations between dominant species across samples. Bubble size indicates p□value, and color represents the correlation coefficient between each pair of SGBs. (**D**) Lollipop chart displaying the coefficient of variation in relative abundance for dominant species. (**E**) Phylogenetic tree constructed from medium- and high-quality bacterial genome assemblies. (**F**) Occurrence frequency of MetaCyc metabolic pathways across bacterial species. (**G**) UMAP clustering of Blautia_A bacterial cells based on genetic features, with colors indicating species identity. (**H**) Heatmap of characteristic UniRef90 protein genes for each species within the genus Blautia_A. (**I**) UMAP clustering of Blautia_A bacterial cells based on genetic features, with colors indicating sample origin.

Applying these criteria, we identified 28,932 high-confidence single cells. The species abundance distribution was characterized by a few dominant taxa—seven species contained over 1,000 cells, led by SGB4933 (4,940 cells) (Top 10 species were showed in **Fig. 4B**). An interaction network among the top 10 species revealed 12 significant pairwise correlations, with a preponderance of negative associations (7 negative vs. 5 positive, **Fig. 4C**), suggesting competitive dynamics may be prevalent in this community. For instance, SGB5121, SGB15318, and SGB15332 formed a cohesive, positively correlated module (Coefficient > 0.5, P < 0.01, **Supplementary Fig. 2**), potentially indicating functional synergy or niche overlap. In contrast, a strong negative correlation was observed between SGB4925 and SGB4262 (Coefficient = 0.71, P = 8.9×10^-5^, **Supplementary Fig. 2**), suggestive of direct competition or antagonism. Notably, one of them, SGB4925, is *Agathobacter faecis*, which has a positive role in the human gut, and its higher abundance is often positively correlated with health status ^20,21^. However, SGB4262 belongs to *Ruminococcus_D bicirculans*, which is usually elevated in unhealthy states ^22,23^, and specific strains were found to be significantly associated with advanced colorectal cancer (CRC), suggesting its potential as a CRC marker^24^. The negative correlation between SGB4925 and SGB4262 may reflect the antagonistic relationship between probiotics and harmful bacteria in the human gut, and also reflect the adaptive regulation of intestinal flora to maintain their own functional stability.

Beyond these specific interactions, we further assessed the overall stability of the microbial community across samples. Community structure was remarkably stable across samples, as evidenced by low coefficients of variation (CV < 0.2) for most species (**Fig. 4D**). However, certain taxa, including SGB4933, SGB1814, and SGB4925, exhibited relatively higher CVs, identifying them as dynamic members potentially responsive to environmental variation. By combining correlation analysis with coefficient of variation information, we aim to identify those species that not only play a key role in community interactions but also have indicative abundance. It is worth noting that for significant negative correlation pairs, such as SGB4925 vs. SGB4262, SGB4925 has a relatively higher CV. This indicates that the competitive relationship between the two species may be mainly driven by the fluctuation of SGB4925, which means that when conditions are suitable for its proliferation, it will effectively inhibit SGB4262. In contrast to the above findings, species that are positively correlated with other members and have extremely low CV (such as SGB5121 and SGB15318) constitute a stable and anti-interference core microbiota. Their stable existence may provide fundamental functions for the entire ecosystem.

#### 2.2 Systematic genomic reconstruction and functional analysis of single-cell genomes

To overcome the limitation of low genomic coverage in individual droplets and reconstruct robust representative genomes, we implemented a co-assembly strategy using the top 50 best-annotated cells from each SGB (for SGBs have less than 10 cells, we use all cells). This approach successfully generated 92 medium- to high-quality metagenome-assembled genomes (MAGs). Taxonomic annotation of these MAGs using GTDB-Tk showed remarkable consistency with the initial SGB classifications derived from MetaPhlAn4 marker genes (**Supplementary Table 2**). Notably, only six SGBs exhibited discrepancies at the species level, but all of these remained concordant at the genus level, with two cases arising from unresolved species annotations in the GTDB database itself. This high concordance validates the reliability of MetaSAG, the marker gene-based classification strategy, for downstream genomic analysis.

A maximum-likelihood phylogenetic tree was then constructed (see **Supplementary Materials and Methods**) from these genomes, capturing substantial evolutionary diversity, encompassing 6 phyla, 9 classes, 15 orders, 23 families, and 59 genera (**Fig. 4E**). The community was dominated by members of the Firmicutes_A phylum (80.43%, 74 genomes), followed by Bacteroidota (11.95%, 11 genomes), reflecting the typical composition of the human gut microbiota.

Subsequently, to reveal the functional capacity of gut microbiota, we annotated metabolic pathways for each species using HUMAnN3 (see **Supplementary Materials and Methods**). As a result, we identified a total of 104 unique MetaCyc pathways. The distribution of these pathways was non-uniform, with each pathway present on average in six species. And 66 pathways were found in more than one species, indicating common functional modules (**Fig. 4F** and **Supplementary Fig. 3**). Among the ten most frequently occurring pathways, four were directly involved in carbohydrate metabolism and synthesis—including sucrose biosynthesis (PWY-7238), glycogen biosynthesis (from ADP−D−Glucose) (GLYCOGENSYNTH-PWY), rhamnose biosynthesis (DTDPRHAMSYN-PWY), and glycolysis IV (PWY-1042)—highlighting the central role of sugar metabolism in gut bacteria.

To dissect the hierarchical association between microbial functional traits and phylogenetic relationships, we assessed the influence of different taxonomic levels (phylum to genus) on functional distribution. Analysis of variance revealed that genus-level classification exerted the most significant effect on functional distribution (Variance = 80.70, Std = 8.98), indicating a pattern of macroscopic functional conservation among closely related species. To investigate whether finer functional distinctions exist among species within the same genus, we further analyzed the composition of marker proteins across species. Clustering analysis based on the presence or absence of UniRef90 protein sequences clearly discriminated between different species within the same genus. Taking the most abundant genus *Blautia_A* as an example, a heatmap based on marker proteins showed that cells robustly clustered by species, forming distinct blocks (**Fig. 4G**). This was quantitatively confirmed by a corresponding hierarchical clustering dendrogram, which demonstrated high intra-species similarity and inter-species dissimilarity (**Fig. 4H**). Notably, this species-specific clustering pattern showed no apparent association with sample origin (**Fig. 4I**), thereby indicating that the functional gene repertoire constitutes a strong species-specific signature that can be robustly captured from single-cell data.

### 3. Analysis of strain level variation and evolution

In bacterial genomics, a species is typically defined as a group of individuals sharing similar genomic characteristics. However, even within the same species, different strains can exhibit significant phenotypic variation, which is often closely associated with genomic single nucleotide polymorphisms (SNPs). Thus, we investigated intra-species heterogeneity by aligning single-cell short-read sequences to assembled species genomes for SNP detection, and then we performed unsupervised clustering on 21 abundant microbial species to identify different strains (see **Supplementary Materials and Methods**).

We observed significant differences in genomic variation frequency across species (**Fig. 5A**). The average SNP distances of SGB4820 and SGB4837 were quite short, with SNPs occurring every 306 and 461 base pairs on average (**Fig. 5A**), which indicated that the genome of this strain is rapidly mutating/accumulating diversity, or its population is rapidly differentiating after experiencing expansion or a bottleneck recently. While SGB4925 has a longer average SNP distance, suggesting that the genomes were nearly identical, with SNPs occurring per 32003 base pairs on average. The extremely low SNP density indicated a recent common ancestor.

**Figure 5.**
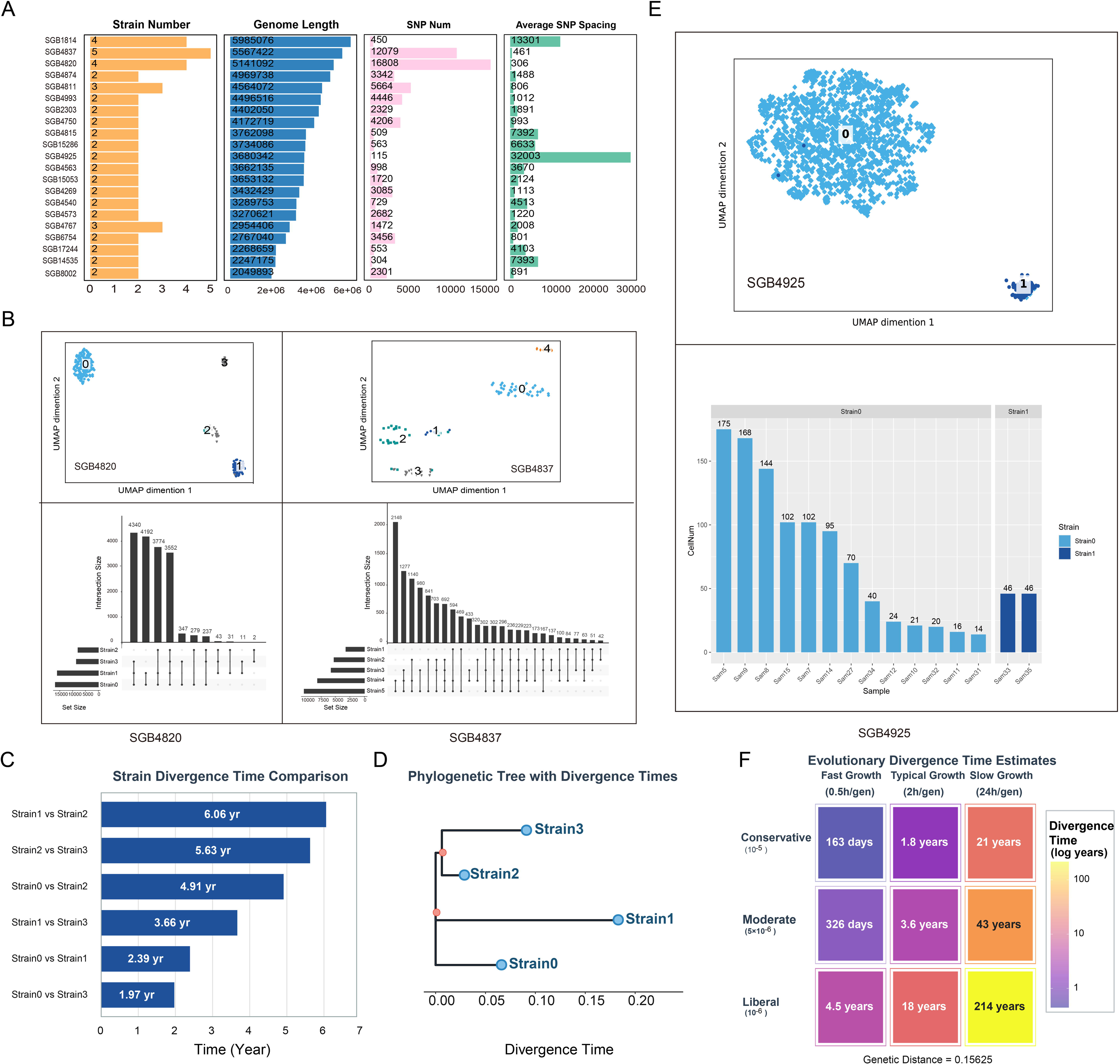
**Analysis of strain-level variation**. (**A**) Statistics of SNP characteristics of strains. (**B**) The UMAP plot of strains and SNPs shared among different strains of SGB4820 and SGB4837. (**C**) The divergence time (year) between strain pairs of SGB4820 under recommended parameters (mutation rate= 5×10^-6^, growth speed= 2h/generation). (**D**) The phylogenetic tree of strains of SGB4820. And the distance between each two strains represents the divergence time. The scale is at the very bottom. (**E**) The clustering of the two strains of SGB4925 and their temporal distribution. (**F**) The evolutionary divergence time estimates of SGB4925 strains under different conditions.

Subsequently, we divided it into different strains through clustering and presented the number of SNP shares among different strains for SGB4820 and SGB4837 (**Fig. 5B**). They respectively have 4 and 5 strains. Taking SGB4820 as an example, we conducted a divergence time estimate using Strain 0 as a reference (**Fig. 5C**, see **Supplementary Materials and Methods**). Under the medium parameter scenario (mutation rate of 5×10^-6^ and generation time of 2 hours), the differentiation times of Strain 1, 2, 3 and Strain 0 were 2.4, 4.9 and 2.0 years, respectively. Based on this information, we constructed a phylogeny tree among the strains (see **Supplementary Materials and Methods**), which can preliminarily infer the evolutionary relationships among different strains (**Fig. 5D**). According to this result, we suggested that they may have differentiated within the same host. We also simulated different scenarios to calculate the differentiation time, and identified top differentiated SNPs that may play a key role in the differentiation of strains in SGB4820 (**Supplementary Fig. 4** and **Supplementary Table 3**).

In addition, SGB4925 consists of two distinct strains (Strain 0 and Strain 1, **Fig. 6E**). Strain 1 emerged abruptly and was detected exclusively in samples collected at the 7th time point (Samples 33 and 35), whereas Strain 0 was detected across the other sampling time points. This pattern suggested a potential colonization or expansion event of Strain 1 between the 6th and 7th sampling points. To investigate their evolutionary relationship, we estimated the divergence time between the two strains. Under the moderate_typical evolutionary scenario, the estimated divergence time was 3.56 years (**Fig. 5F** and **Supplementary Table 4**). Notably, the actual interval between the sixth and seventh sampling points was 121 days (approximately 0.33 years), which aligns with the conservation_fast parameter scenario. Overall, most evolutionary scenarios supported a recent divergence of the two strains.

**Figure 6.**
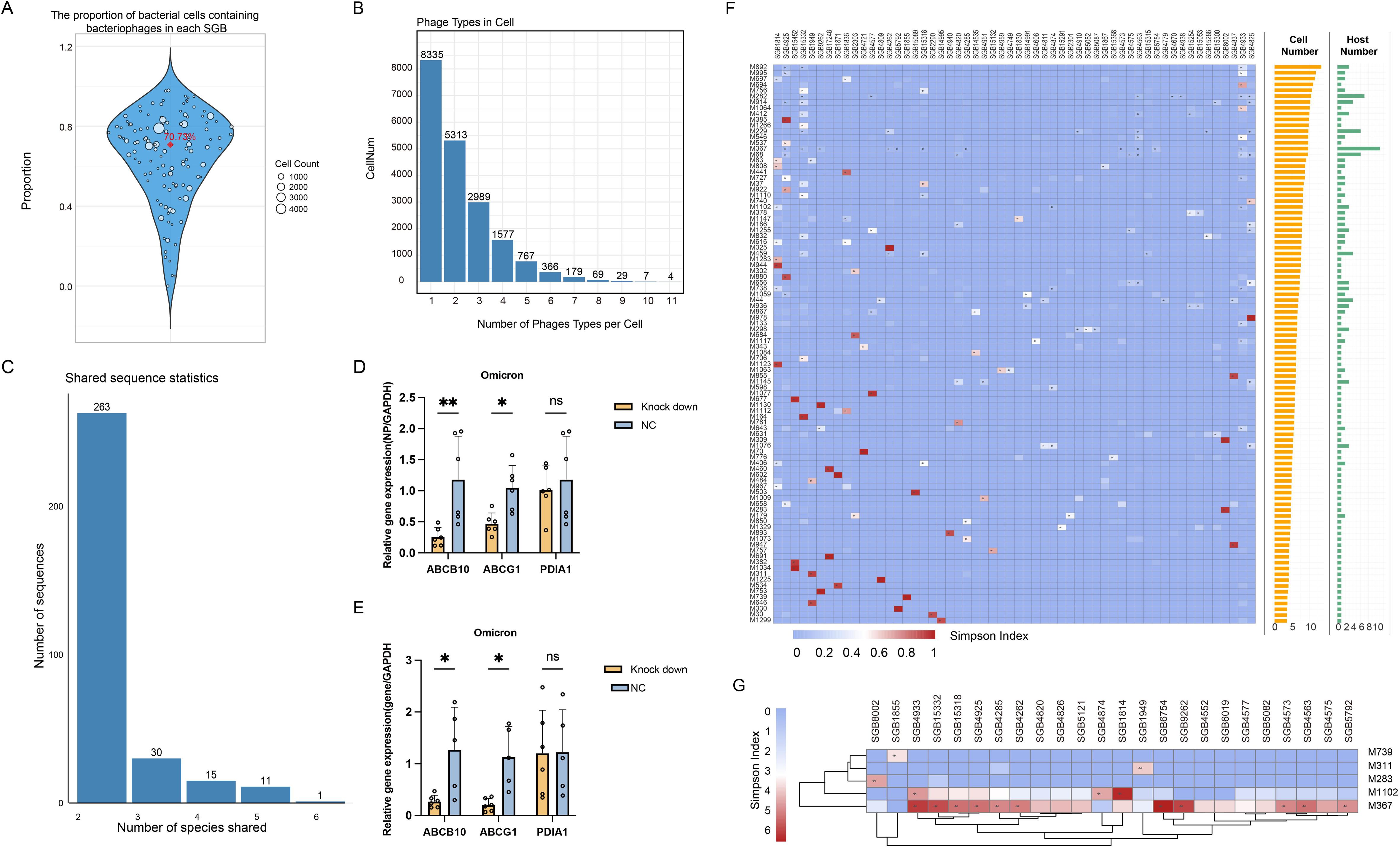
Interaction between bacteriophages and bacteria. (**A**) Violin plot of the proportion distribution of phage infected droplets in each bacterial species. Each bubble represents a species, and the size means the proportion of the bacteria infected by phages. (**B**) The number of bacteria cells that infected by one or more types of phages. (**C**) The statistic of the sequence count shared by multiple species. (**D**) Viral NP transcript level (NP/GAPDH). **(E)** Target gene transcript level (gene/GAPDH). (**F**) The infection relationship between bacteria and phages. The color represents the Simpson index, and the symbol ‘*’ means that the relationship between them is significant. (**G**). The infection relationship between three specific phages and two broad-spectrum phages and bacteria.

To understand the genetic basis of the strain differentiation of SGB4925, we functionally annotated the SNPs. Remarkably, approximately 33% (50/151) of the SNPs were clustered within transposase genes and associated insertion sequence (IS) elements, primarily from the IS116/IS110/IS902 and IS66 families (**Supplementary Table 4**). This implicated that mobile genetic element may be a key driver of genetic instability and strain diversification in SGB4925. Among the 30 non-synonymous mutations identified, several were predicted to have significant functional consequences: (i) The R37S mutation in an IS66 family transposase, which substitutes a basic for a polar neutral residue, potentially disrupting DNA-binding affinity or catalytic activity; (ii) The E106K mutation, also within the IS66 transposase, which replaces an acidic residue with a basic one. As this residue is located within the conserved DDE motif critical for Mg²□ coordination, this mutation is likely to abolish transposition activity entirely; (3) The D76H mutation in an IS66 C-terminal element, causing a charge reversal that could alter protein-DNA or protein-protein electrostatic interactions. These findings highlight that transposition-related mutations, particularly those impacting catalytic cores and DNA-binding interfaces, may underpin the functional divergence and dynamic behavior of the strain types of SGB4925.

### 4. A Single-Cell Atlas of Phage-Host Interactions Unveils Susceptible Bacteria and Broad-Host-Range Phages

#### 4.1 Identification of susceptible bacterial hosts as key targets

The encapsulation of individual microbes provides an ideal framework to study phage-host relationships without the averaging effect of bulk sequencing. This is of great significance for identifying lytic phages and facilitating the research of phage therapy. To discover the invasive relationship between phages and bacteria, we first established a confidence threshold for phage detection, excluding phage annotations with fewer than 80 assigned reads per cell, as validation with simulated data indicated these low-abundance signals were likely stochastic (**Supplementary Fig. 5**, see **Supplementary Materials and Methods**). As a result, phages were found to be widespread symbionts of gut bacteria: the median percentage of phage-positive cells within a bacterial species was 70.73% (**Fig. 6A**), and the number of infected cells per species was strongly correlated with its total cell count (**Supplementary Fig. 6A**), suggesting that phage abundance is tightly linked to host availability. Besides, over half of the phage-positive cells carried two or more distinct phage types, demonstrating that co-occurrence within single bacterial cells was common (**Fig. 6B**). One extreme case, Cell505945 (identified as *Faecalibacterium prausnitzii*, SGB15332), harbored 11 different phages (**Fig. 6B**), illustrating the potential for complex viral consortia within a single host and that this kind of bacteria may be easily infected by bacteriophages.

To preliminarily investigate the potential mechanisms underlying the susceptibility of these bacterial hosts to phage infection, we first identified susceptible bacteria. For each bacterial species, the proportion of cells harboring phage (PhageBac_CellsProp) was calculated. Additionally, the Shannon diversity index was also computed based on the read counts of different phages present within each bacterial species, with higher values indicating a greater number of phage species and a more even distribution. Using these metrics, we selected nine highly susceptible bacterial species (criteria: PhageBac_CellsProp > 0.7, Shannon diversity index > 2, and Genome_quality = High; see **Supplementary Table 5**). We obtained the shared amino acid sequences (conserved amino acid sequences) among these nine species for initial screening of targets that may play a role in phage-host recognition. Notably, one sequence was shared across six SGBs, and eleven sequences were shared among five SGBs (**Fig. 6C, Supplementary Table 5**).

The identification of these shared sequences represents a key finding of our study, as they pinpoint a minimal set of high-priority candidate factors that could underlie broad phage-host interaction ranges. For example, through blastp comparison, we identified that a certain sequence shared by five susceptible bacteria (S3, **Supplementary Table 5**) overlapped with the DUF6551 family protein, which is an unknown functional domain frequently found in phages. This suggests that this sequence might have been passed from a phage to the bacteria^25^. Furthermore, in another sequence (S4), we found that it was aligned with the MbpC protein, on which there was a transmembrane region (K9DVT2_9BACE, position: 100-123), suggesting that this sequence might be related to signal transduction and could be one of the potential ways for phages to infect bacteria.

#### 4.2 Validation of susceptible genes in viral infection

To validate the functional relevance of susceptible genes identified from bacterial protein sequences, we mapped it to the homologous sequence of humans and selected the top three susceptibility genes (ABCB10, ABCG1, and PDIA1) for viral infection experiments. We performed siRNA knockdown of ABCB10, ABCG1, and PDIA1 in A549-TMPRSS2-ACE2 cells followed by SARS-CoV-2 Omicron infection **(Fig. 6D, E)**. The results showed that knockdown of ABCB10 significantly reduced viral NP transcript levels compared with the negative control (NC) (P < 0.01), indicating that ABCB10 promotes Omicron infection.

Similarly, ABCG1 knockdown led to a significant decrease in viral NP expression (P < 0.05), suggesting that ABCG1 is also required for efficient viral replication. In contrast, PDIA1 knockdown did not significantly alter viral NP levels (P > 0.05), implying that PDIA1 may not be essential for Omicron infection under these conditions. RT–qPCR confirmed effective knockdown of ABCB10 and ABCG1 (P < 0.05), whereas PDIA1 knockdown efficiency did not reach statistical significance (P > 0.05) **(Fig. 6D,E)**. Furthermore, transcriptomic analysis of publicly available SARS-CoV-2 datasets revealed that ABCB10 and ABCG1 were frequently upregulated across multiple GEO datasets **(Supplementary Fig. 6B)**, consistent with a potential proviral role. These results indicate that ABCB10 and ABCG1, but not PDIA1, contribute to Omicron infection in A549-TMPRSS2-ACE2 cells and may represent potential host factors for therapeutic targeting.

#### 4.3 Characterization of broad-host-range phages

Given the wide distribution of phages, we also explored their host specificity or broad-spectrum nature at the single-cell level. Phages that can only infect a single type of bacterial host are defined as host-specific phages, and those that can infect multiple hosts are defined as broad-spectrum phages. To distinguish between the two, we used the Simpson index to quantify the parasitic diversity of phages on bacterial hosts and statistically analyzed the phage-host relationship within the range that contributed more than 90% of the host diversity (**Supplementary Fig. 6C, see Supplementary Materials and Methods**). This analysis identified 58 phages with a highly specific tropism, infecting only a single bacterial host (**Fig. 6F**). Among these were three previously characterized phages—M739 (infecting *Bacteroides fragilis*), M283 (infecting *Streptococcus thermophilus*), and M311 (infecting *Parabacteroides merdae*)—whose documented host specificities were perfectly recapitulated in our data, serving as a positive control for our network inference (**Fig. 6G**).

Conversely, we also discovered numerous generalist phages capable of infecting multiple bacterial species (**Fig. 6G**). The most promiscuous phage, a known enterobacterial phage M367 (taxid 414970), was detected in 11 different bacterial species, with its most frequent hosts being SGB6754 (*Faecalibacillus intestinalis*) and SGB4933 (*Agathobacter rectalis*). It is worth noting that among the 11 types of bacteria infected by M367, 10 belong to the same phylum (*Firmicutes_A*), and 8 of them belong to *Clostridia* class, including two orders: *Lachnospirales* (4) and *Oscillospirales* (4). This indicates that bacteria with kinship are more likely to be infected by the same type of phage. In the same way, the consensus features of these phages (M367 and M1102) were preliminarily analyzed to provide researchers with candidate targets for the development of phage therapy (**Supplementary Table 6, 7**).

### 5. MetaK-Lytic: A Few-Shot Meta-Learning Framework for Predicting Phage Lytic Ability from short reads

We have identified the parasitic relationship between phages and their hosts, but the nature of these phages remains unknown, such as whether they cause host lysis. However, since most phages are uncultured, such information about them is difficult to obtain. Therefore, an algorithm is needed to predict their lytic ability. Conventional identification methods that rely on marker genes are often infeasible for SAG data due to incomplete genome recovery. This creates a central challenge: how to determining phage lytic ability in the absence of known marker genes or large labeled datasets? To address this, we developed MetaK-Lytic, a few-shot meta-learning framework that directly based on k-mer genomic sequence features to infer phage lytic ability (**Fig. 7A**).

**Figure 7.**
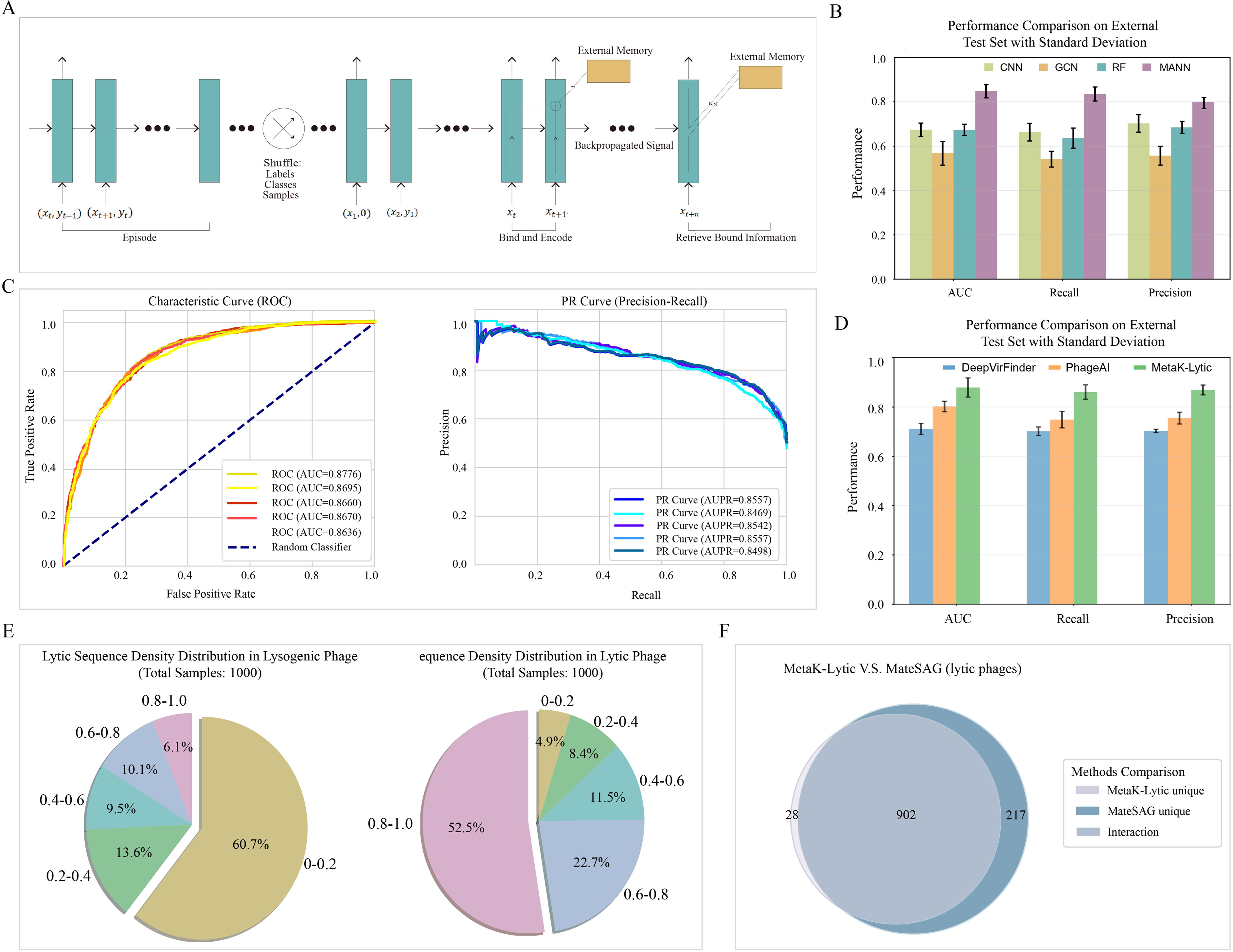
Development and performance demonstration of MetaK-Lytic. (**A**) The development process of MetaK-Lytic. (**B**) The performance of multiple algorithms compared to MANN in AUC, recall, and precision. (**C**) The ROC and PR curves from five randomized trials by MetaK-Lytic using the same data. (**D**) The performance comparison of MetaK-Lytic and other existing methods on external test set. (**E**) Density distribution of predicted lytic sequences in lysogenic phages and lytic phages. (**F**) Analysis of concordance between phages predicted to be lytic by MetaK-Lytic and those identified as lytic by MetaSAG.

**Figure 8.** Effects of ABCB10, ABCG1 and PDIA1 knockdown on Omicron infection and gene expression. Cells transfected with ABCB10/ABCG1/PDIA1 knockdown reagents or negative control (NC) were subjected to Omicron treatment/infection. RT–qPCR was used to quantify relative transcript levels, normalized to GAPDH. **(A)** Viral NP transcript level (NP/GAPDH). **(B)** Target gene transcript level (gene/GAPDH). **(C)** Heatmap of ABCB10 and ABCG1 expression changes across SARS-CoV-2 transcriptomic datasets. Color intensity indicates log2 fold change (log2FC), with white for zero and red for higher values.

We first built a feature-extraction pipeline that converts obtained phage genomes (training data) into numerical representations suitable for meta-learning (see **Supplementary Materials and Methods**). Complete genomes were fragmented in silico to simulate short-read sequencing, from which we computed 31-mer frequency spectra as alignment-free sequence fingerprints. It showed that 31-mers offer sufficient sequence specificity while remaining computationally tractable. The resulting feature vectors capture global sequence composition without requiring gene annotation.

The core of MetaK-Lytic is a MANN architecture equipped with a Least-Recently-Used Access (LRUA) memory module, enabling rapid storage and retrieval of task-specific information. During training, the model was optimized in an episode-based regime: each episode presented a randomly sampled few-shot classification task, and class labels were permuted across episodes. This design forces the model to rely on its external memory rather than memorizing fixed label associations (**Fig. 7A**, see **Supplementary Materials and Methods**).

We benchmarked MANN against several baseline algorithms, including convolutional neural network (CNN), graph convolutional network (GCN), and random forest (RF). MANN outperformed all baselines, achieving 85.3% accuracy for the sequence prediction in a five-shot setting on an independent external dataset (**Fig. 7B**), demonstrating strong generalization from very few labeled examples.

To evaluate stability, we performed five replicate runs of MetaK-Lytic on the same data for predicting labels of short sequences. Results showed consistent and excellent performance, with AUC values consistently exceeding 85%, as well as high precision and recall (**Fig. 7C**). We also compared MetaK-Lytic against existing predictors based on marker gene ^18^ and long-read sequences ^26^. Our method showed superior performance, improving accuracy by more than 10% over other approaches and reaching up to 95% accuracy (**Fig. 7D**).

We then applied MetaK-Lytic to the comprehensive catalog of phages identified from our SAG data. Each fragment within a phage genome was assigned a lytic or non-lytic label. Based on the probability distribution of these fragments (**Fig. 7E** and **Supplementary Fig. 7A**), we defined a threshold (0.8) to characterize whole phages: a phage containing >80% lytic fragments was classified as lytic.

For further validation, we used intracellular phage-read abundance to indicate lytic activity. By analyzing the distribution of phage-read proportions across all infected cells, we defined a putative lytic-state threshold of 0.1715 (>17.15% of cellular reads mapping to a specific phage; **Supplementary Fig. 7B, C**). These results were consistent with MetaK-Lytic predictions: 97% of phages identified as lytic by MetaK-Lytic were also lytic according to the read-abundance criterion (**Fig. 7F** and **Supplementary Table 8**).

In summary, MetaK-Lytic is a short-sequence-based approach that does not rely on marker-gene libraries, can be applied to uncultured phages, and is highly suited to microbial single-cell genome sequencing data. This framework provides a powerful tool for in silico assessment of phage lytic ability and can help accelerate the selection of candidate phages for therapeutic development.

### 6. MetaSAG: An Integrated Software Suite for Single-Cell Microbial Sequencing Data Analysis

To translate our methodological advances into a practical resource, we have integrated all analytical components into MetaSAG, an open-source software suite (https://github.com/liangcheng-hrbmu/MetaSAG). MetaSAG provides a modular, reproducible workflow that transforms raw single-cell sequencing data into interpretable results through the following core modules (**Fig. 1**).

#### 6.1. Adaptive Data Preprocessing and Quality Control

MetaSAG begins with data-driven quality control, automatically determining the optimal read-count cutoff by identifying the inflection point in the droplet read distribution, thereby maximizing data retention while removing low-quality droplets (**Fig. 2A-C**).

#### 6.2. High-Confidence Taxonomic Profiling

Using the MetaPhlAn4 marker database, this module rapidly assigns species-level identities to each droplet, outperforming general classifiers in accuracy, and reports potential co-encapsulation events (**Fig. 2** and **Fig. 3**).

#### 6.3. Functional Profiling and Phylogenetic Analysis

Cells of the same species are co-assembled into metagenome-assembled genomes (MAGs). MetaSAG then reconstructs phylogenetic trees and profiles functional potential with HUMAnN3, linking taxonomy to metabolic capacity (**Fig. 4**). Besides, the horizontal gene transfer (HGT) analysis is provided, which is able to identify gene transfer events between different species (**Supplementary Fig. 8, Supplementary Table 9**).

#### 6.4. Strain-Level Variation and Dynamics

By detecting SNPs from single-cell reads aligned to MAGs, MetaSAG clusters cells into strains and quantifies intra-species diversity. For longitudinal data, it can track strain emergence and turnover over time (**Fig. 5**).

#### 6.5. Phage-Host Interaction Mapping and Lyticity Prediction

This module builds single-cell resolution phage–host networks and integrates **MetaK-Lytic**—our few-shot learning framework that predicts lytic vs. temperate lifestyle directly from k-mer profiles, without marker genes (**Fig. 6, 7**).

All modules are orchestrated in a single workflow. MetaSAG accepts standard FASTQ inputs and generates interpretable reports. The tool is freely available under the MIT License at: https://github.com/liangcheng-hrbmu/MetaSAG.

## Discussion

This study developed the MetaSAG platform, providing a comprehensive solution for microbial single-amplified genome data analysis, spanning from raw sequencing reads to strain evolution, community structure, and phage-host interactions. The platform demonstrates notable advantages in several aspects and holds broad significance both methodologically and in application.

First, MetaSAG achieves important breakthroughs in the analytical workflow. We introduced a data-driven droplet quality control strategy that adaptively identifies and retains high-quality single-cell data, significantly enhancing the reliability of downstream analyses. In terms of taxonomic classification, MetaSAG employs the MetaPhlAn4 marker gene database as its core classifier, maintaining high classification confidence while substantially improving processing speed. Compared to traditional co-assembly methods, computational time is reduced by approximately 87.9%, making large-scale single-cell microbiome studies feasible on conventional computing platforms.

Second, MetaSAG not only achieves precise species-level identification but also extends analysis to the strain level for evolutionary insights. At single-cell resolution, we systematically characterized the strain diversity of human gut microbiota and preliminarily revealed the dynamic succession patterns and potential evolutionary timelines of certain strains.

Furthermore, by integrating phage detection and host association analysis, MetaSAG successfully constructed a single-cell resolution bacterial-virus interaction network, identifying multiple broad-host-range phages and susceptible bacterial hosts. This provides crucial leads for subsequent phage resource mining and interaction mechanism studies.

More importantly, the MetaSAG platform offers a practical tool for addressing long-standing technical challenges such as exploring microbial “dark matter,” detecting low-abundance taxa, and analyzing functional heterogeneity at the strain level. In basic research, the platform can be used to deeply investigate microbial community assembly mechanisms, species interaction networks, and microevolutionary processes. In clinical applications, MetaSAG can track pathogen transmission chains, identify disease-associated functional strains or antibiotic-resistant variants, and screen for phages with therapeutic potential. This provides new perspectives and data support for infectious disease control and the development of microbiome-based therapies.

However, this study has several limitations to be further explored. First, although we performed strain typing and evolutionary inference based on SNPs, we did not deeply explore the functional implications of these SNPs. The identified genetic variations were mostly concentrated in regions related to transposases, and whether these mutations truly affect bacterial adaptability or pathogenicity requires experimental validation. Second, our analysis of phage-host interactions remains at the correlative level, and the molecular mechanisms have not been thoroughly investigated. Although we identified a set of conserved sequences potentially involved in interactions, their specific functions need further confirmation through structural biology or biochemical experiments. Additionally, MetaSAG relies on threshold-based strategies for identifying co-encapsulation events, which may miss signals from low-abundance or highly similar co-encapsulated species. Future studies could integrate long-read sequencing or optical mapping technologies to improve detection sensitivity. Finally, our characterization of the dynamic co-evolution between phages and hosts remains preliminary. Applying this workflow to more densely sampled time-series data in the future may allow clearer elucidation of virus-host co-evolutionary trajectories.

In summary, MetaSAG represents an efficient, multi-level single-cell microbiome analysis platform that holds promising application prospects. We look forward to further advancing the field of microbiomics toward higher resolution and functional interpretability through continuous methodological refinement and deeper integration with experimental research.

## Materials and methods

### 1. Microbe-seq Data Processing

Microbe-seq libraries were prepared as previously described ^9^. In brief, they isolate microbes by encapsulating them into droplets with lysis reagents using a microfluidic device. The droplet emulsion is then transferred to a syringe and reinjected into a droplet generator to form droplets encapsulating Multiple Displacement Amplification (MDA) reagents, thereby enabling the cultivation and amplification of microbial genomes.

Raw sequencing data were processed using Trimmomatic (version 0.36) with the following parameters: [*ILLUMINACLIP=’TruSeq3-PE.fa:2:30:10:3:TRUE’, LEADING=25, TRAILING=3, SLIDINGWINDOW=’4:20’, MINLEN=30*] to remove adapters and low-quality reads.

### 2. Droplet Quality Control and Read Count Filtering

The read count per droplet (*r_d_*) was calculated. The optimal threshold (*r\**) for filtering low-quality droplets was determined by identifying the inflection point of the empirical cumulative distribution function (ECDF) of *r_d_*.

The ECDF was computed using the ecdf() function in R (version 4.2.2) and visualized using R package “ggplot2”. The inflection point was identified by numerically approximating the second derivative using the finite difference method and locating the point where it crossed zero. Droplets with *r_d_* < *r\** were filtered out.

### 3. Taxonomic Annotation with MetaSAG and Comparative Tools

The MetaSAG pipeline was implemented, utilizing the MetaPhlAn4 database as its core classifier. The annotation was performed with default parameters.

For comparative analysis, Kraken2 and Kaiju were also used under standard parameters: [*Kraken2: --confidence 0.0,--minimum-base-quality 0,--minimum-hit-groups 2; Kaiju: -a greedy -e 3 -m 11 -s 65 -E 0.01*].

Only species-level classifications were retained.

### 4. Validation of Species-Level Classifications

**Metagenomic-Based Validation**: Genomes from individual droplets were assembled using SPAdes (version 3.13.0) with parameters: [--sc –careful]. Contigs from all droplets were pooled, and ribosomal sequences were identified and annotated using QIIME2 (version 2023.2.0) against the SILVA 138 database.

**Genome Similarity-Based Validation**: Following the methodology of Zheng et al. ^9^, cells were clustered based on genomic similarity. In brief, filtering, individual assembly with SPAdes, genomic similarity calculation with Sourmash, iterative co-assembly, and final clustering bins at ANI ≥95%. The resulting clusters were annotated using GTDB-Tk (version 1.0.2).

### 5. Identification of susceptible genes from bacterial protein sequences

Susceptible genes were identified through a multi-step screening pipeline. First, protein sequences that were present in at least 3 susceptible bacterial species were retained. These sequences were then used as queries for protein BLAST (blastp) against the NCBI non-redundant protein database, with the search restricted to human proteins (taxid: 9606). For each query sequence, the human protein hit with the smallest E-value was selected as the best match. The matched protein was subsequently mapped to its corresponding human gene to obtain the list of susceptible genes.

### 6. Gene knockdown experiments

**Cells and viruses**: SARS-CoV-2 Omicron BA.5 (GenBank: OM21247) was used in this study, virus propagation was conducted as previously described ^27^. Virus stocks were titrated by plaque assay, aliquoted, and stored at −80 °C. A549-TMPRSS2-ACE2 cells were maintained in DMEM supplemented with 10% fetal bovine serum (FBS), 1% penicillin–streptomycin (P/S), 0.5 μg/mL puromycin, and 300 μg/mL hygromycin for selection. All experiments involving live SARS-CoV-2 were conducted in accordance with approved standard operating procedures in the biosafety level 2 facility at the University of Hong Kong.

**Transfection and siRNA screening**: siRNAs targeting the mRNA of selected gene candidates were obtained from Thermo Fisher Scientific or synthesized by Tsingke Biotech (Beijing, China). A nontargeting (scrambled) siRNA from Tsingke Biotech served as the negative control. For siRNA transfection, A549-TMPRSS2-ACE2 cells were seeded in 48-well plates at 40,000 cells per well and transfected with a 20 μL mixture containing 5 pmol siRNA and 1.5 μL Lipofectamine™ RNAiMAX Transfection Reagent (Thermo Fisher Scientific) in Opti-MEM reduced serum medium, according to the manufacturer’s instructions. At 24 h post-transfection, the medium was removed and cells were infected with SARS-CoV-2 at a multiplicity of infection (MOI) of 0.01 for 48 h. Cell lysates were prepared using RLT lysis buffer, and total RNA was extracted with the RNeasy kit (Qiagen, Hilden, Germany). Viral and target gene transcript levels were quantified by one-step RT–qPCR using the TB Green® PrimeScript™ RT-qPCR Kit II (Takara, Kusatsu, Japan) with SARS-CoV-2 NP and target gene-specific primers, as described previously ^27^. Gene expression was normalized to GAPDH.

### 7. Statistical Analysis and Visualization

All statistical analyses were performed in R (version 4.3.1). Visualization of phylogenetic trees was performed using the ape and ggtree packages.

**Quantitative analysis of gene expression**: The heatmap displays log2 fold changes (log_2_FC) of ABCB10 and ABCG1 across SARS-CoV-2-related transcriptomic datasets. Gene expression data were downloaded from the Gene Expression Omnibus (GEO) (https://www.ncbi.nlm.nih.gov/geo/). Differential expression analysis was performed by comparing SARS-CoV-2 infection with mock. The count-based high-throughput sequencing data for the transcriptome was standardized to Transcripts Per Million (TPM). Subsequently, the TPM data, along with other high-throughput sequencing data formats such as FPKM and RPKM, underwent normalization using the limma package. Following normalization, a logarithmic transformation with log2 was applied to all transcriptomic data. Datasets were selected if at least one of the two genes showed significant upregulation (log_2_FC ≥ 1 and P < 0.05). Statistical significance was indicated by asterisks (* P < 0.05, ** P < 0.01, *** P < 0.001, **** P < 0.0001) overlaid on each cell.

## Data and Code Availability

The combined fastq files have been deposited in the NCBI Sequence Read Archive under the BioProject accession number PRJNA803937. And the Metagenomic fastq files are available from the previous publication (Bioproject: PRJNA544527).

The MetaSAG pipeline, including the MetaK-Lytic model are open-source and publicly available at https://github.com/liangcheng-hrbmu/MetaSAG.

## Acknowledgments

This work was supported by the National Natural Science Foundation of China (62222104, 62531007, and 62172130), and Heilongjiang Postdoctoral Fund (LBH-Q20030, LBH-Z24238).

## Author Contributions

Liang Cheng and Xue Zhang conceived and headed the project. Hongbin Han participated in the design of the project and the revision of the manuscript. Sainan Zhang and Meiyu Du designed and participated in the raw data processing and the technical analysis. Guanzhi He designed and developed the algorithm (MetaK-Lytic). Kai Qian, Baifeng Li, Ping Wang, Xiaoliang Wu, and Chao Wang contributed to the improvement of the tool. Shuofeng Yuan and Kexin Li designed and performed the biological experiment. Sainan Zhang drafted the manuscript. Minke Lu and Liang Cheng revised the manuscript. All authors approved the final version of the manuscript.

## Competing Interest Statement

The authors declare no competing financial interests.

## Notes

https://github.com/liangcheng-hrbmu/MetaSAG

### Competing Interest Statement

The authors have declared no competing interest.

## References

1. Wastyk, H.C., et al. Gut-microbiota-targeted diets modulate human immune status. Cell 184, 4137–4153 e4114 (2021).

2. Basolo, A., et al. Effects of underfeeding and oral vancomycin on gut microbiome and nutrient absorption in humans. Nat Med 26, 589–598 (2020).

3. Kayama, H., Okumura, R. & Takeda, K. Interaction Between the Microbiota, Epithelia, and Immune Cells in the Intestine. Annu Rev Immunol 38, 23–48 (2020).

4. Rosselli, R., et al. Direct 16S rRNA-seq from bacterial communities: a PCR-independent approach to simultaneously assess microbial diversity and functional activity potential of each taxon. Sci Rep 6, 32165 (2016).

5. Braukmann, T.W.A., et al. Metabarcoding a diverse arthropod mock community. Mol Ecol Resour 19, 711–727 (2019).

6. Allen, J.P., Snitkin, E., Pincus, N.B. & Hauser, A.R. Forest and Trees: Exploring Bacterial Virulence with Genome-wide Association Studies and Machine Learning. Trends Microbiol 29, 621–633 (2021).

7. Ikhimiukor, O.O., et al. Long-term persistence of diverse clones shapes the transmission landscape of invasive Listeria monocytogenes. Genome Med 16, 109 (2024).

8. Lee, J.J., et al. Trehalose catalytic shift inherently enhances phenotypic heterogeneity and multidrug resistance in Mycobacterium tuberculosis. Nat Commun 16, 6442 (2025).

9. Zheng, W., et al. High-throughput, single-microbe genomics with strain resolution, applied to a human gut microbiome. Science 376, eabm1483 (2022).

10. Bokulich, N.A., et al. Optimizing taxonomic classification of marker-gene amplicon sequences with QIIME 2’s q2-feature-classifier plugin. Microbiome 6, 90 (2018).

11. Wood, D.E., Lu, J. & Langmead, B. Improved metagenomic analysis with Kraken 2. Genome Biol 20, 257 (2019).

12. Menzel, P., Ng, K.L. & Krogh, A. Fast and sensitive taxonomic classification for metagenomics with Kaiju. Nat Commun 7, 11257 (2016).

13. Blanco-Miguez, A., et al. Extending and improving metagenomic taxonomic profiling with uncharacterized species using MetaPhlAn 4. Nat Biotechnol 41, 1633–1644 (2023).

14. Vatanen, T., et al. Genomic variation and strain-specific functional adaptation in the human gut microbiome during early life. Nat Microbiol 4, 470–479 (2019).

15. Dedrick, R.M., et al. Engineered bacteriophages for treatment of a patient with a disseminated drug-resistant Mycobacterium abscessus. Nat Med 25, 730–733 (2019).

16. Kunisch, F., et al. Targeting Pseudomonas aeruginosa biofilm with an evolutionary trained bacteriophage cocktail exploiting phage resistance trade-offs. Nat Commun 15, 8572 (2024).

17. Jault, P., et al. Efficacy and tolerability of a cocktail of bacteriophages to treat burn wounds infected by Pseudomonas aeruginosa (PhagoBurn): a randomised, controlled, double-blind phase 1/2 trial. Lancet Infect Dis 19, 35–45 (2019).

18. Ren, J., et al. Identifying viruses from metagenomic data using deep learning. Quant Biol 8, 64–77 (2020).

19. McNair, K., Bailey, B.A. & Edwards, R.A. PHACTS, a computational approach to classifying the lifestyle of phages. Bioinformatics 28, 614–618 (2012).

20. Zakerska-Banaszak, O., et al. New potential biomarkers of ulcerative colitis and disease course - integrated metagenomic and metabolomic analysis among Polish patients. J Gastroenterol 60, 1384–1399 (2025).

21. Zhang, L., et al. The Intestinal Microbiota Composition in Early and Late Stages of Diabetic Kidney Disease. Microbiol Spectr 11, e0038223 (2023).

22. Noel, S., et al. Metagenomic Profiling of Gut Microbiota in Kidney Precision Medicine Project Participants With CKD and AKI. Compr Physiol 15, e70058 (2025).

23. Wu, Y., et al. Aging characteristics of colorectal cancer based on gut microbiota. Cancer Med 12, 17822–17834 (2023).

24. Piccinno, G., et al. Pooled analysis of 3,741 stool metagenomes from 18 cohorts for cross-stage and strain-level reproducible microbial biomarkers of colorectal cancer. Nat Med 31, 2416–2429 (2025).

25. Kaljevic, J., Sukhoverkov, K.V., Johnson, K.E., Hocher, A. & Le, T.B.K. Versatile NTP recognition and domain fusions expand the functional repertoire of the ParB-CTPase fold beyond chromosome segregation. Proc Natl Acad Sci U S A 122, e2527592122 (2025).

26. Tynecki, P., et al. PhageAI - Bacteriophage Life Cycle Recognition with Machine Learning and Natural Language Processing. bioRxiv, 2020.2007.2011.198606 (2020).

27. Yuan, S., et al. Clofazimine broadly inhibits coronaviruses including SARS-CoV-2. Nature 593, 418–423 (2021).

