## Supplementary Figures for "MetaSAG: A Tool for Multi-level Exploration and Taxonomic Analysis of Microbial Single-Amplified Genomes"

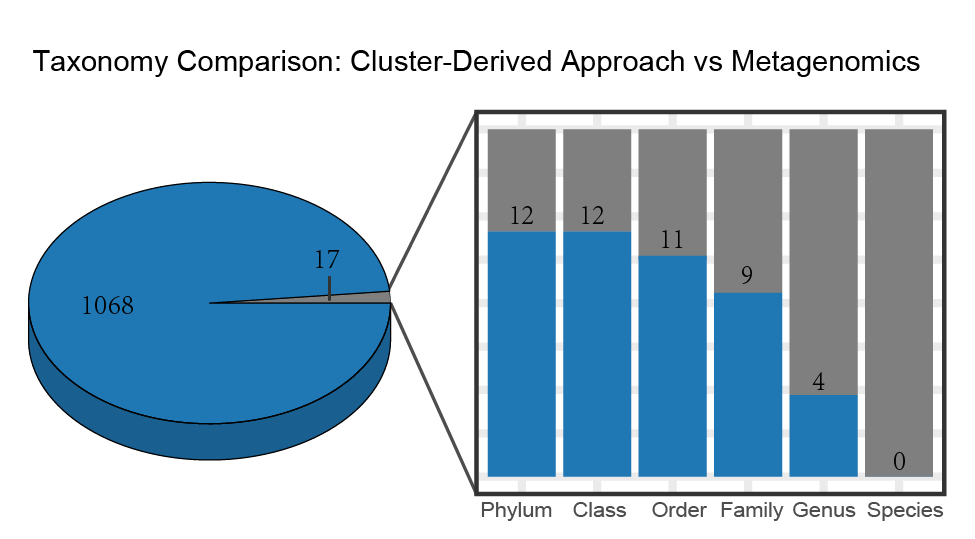

**Supplementary Figure 1. The consistent and inconsistent species that Metagenomics intersected with Cluster-Derived Approach and Metagenomics.** Blue indicates agreement and gray indicates disagreement. The bar graph represents the distribution of agreement at each taxonomic level for species with inconsistent classification.

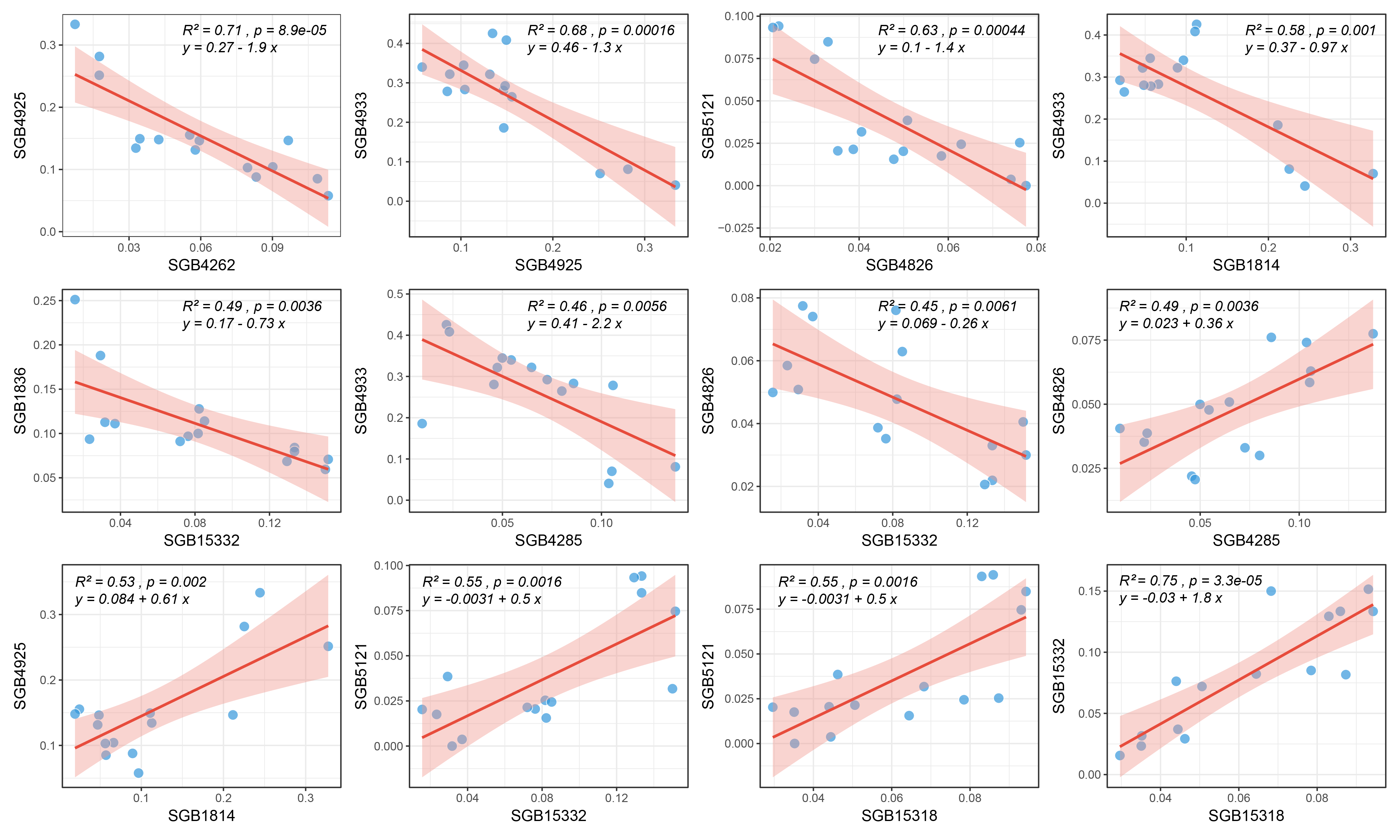
**Supplementary Figure 2. The correlation of dominant species.**

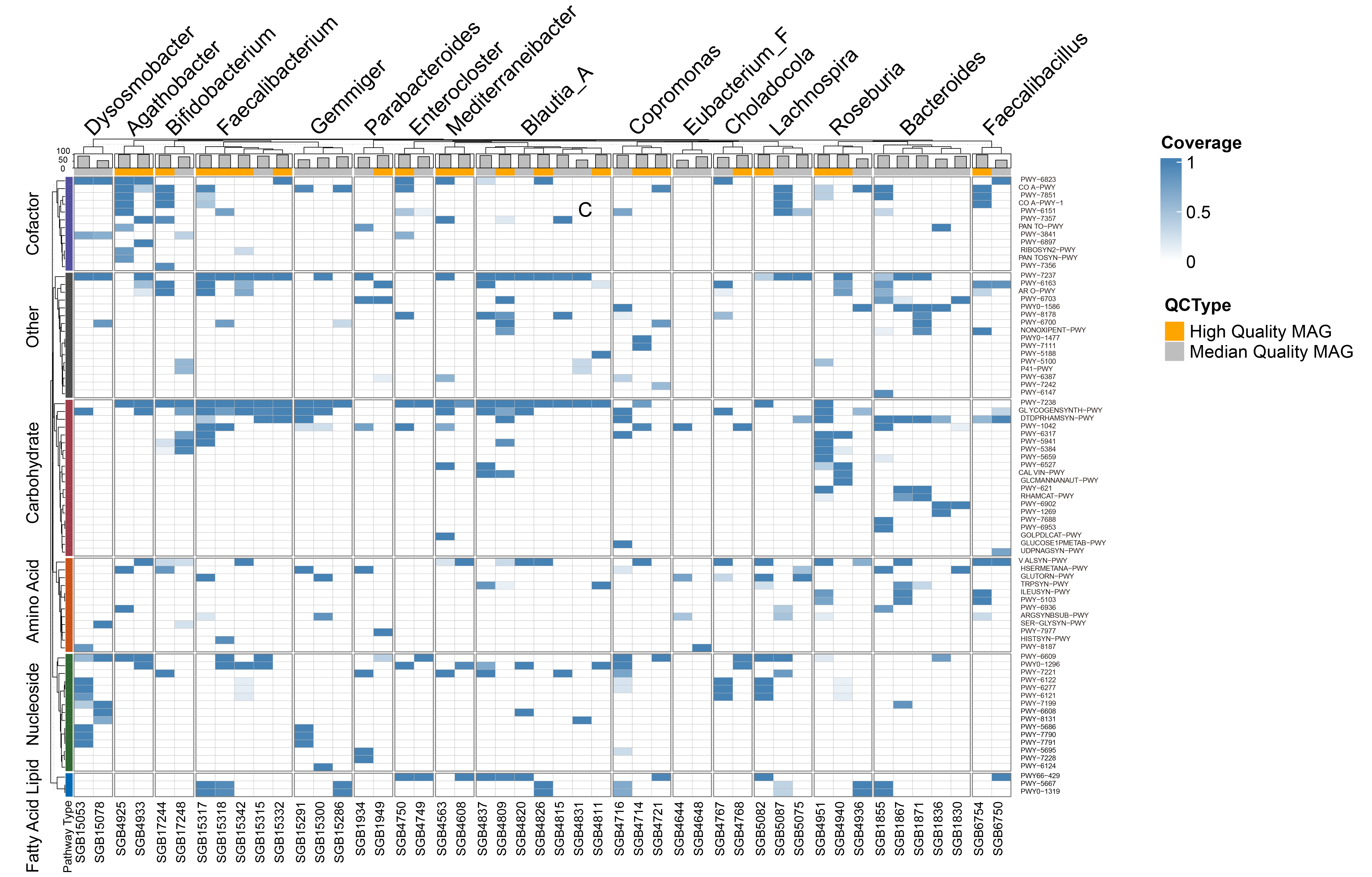

**Supplementary Figure 3. Enrichment of SGBs in multiple pathways.**

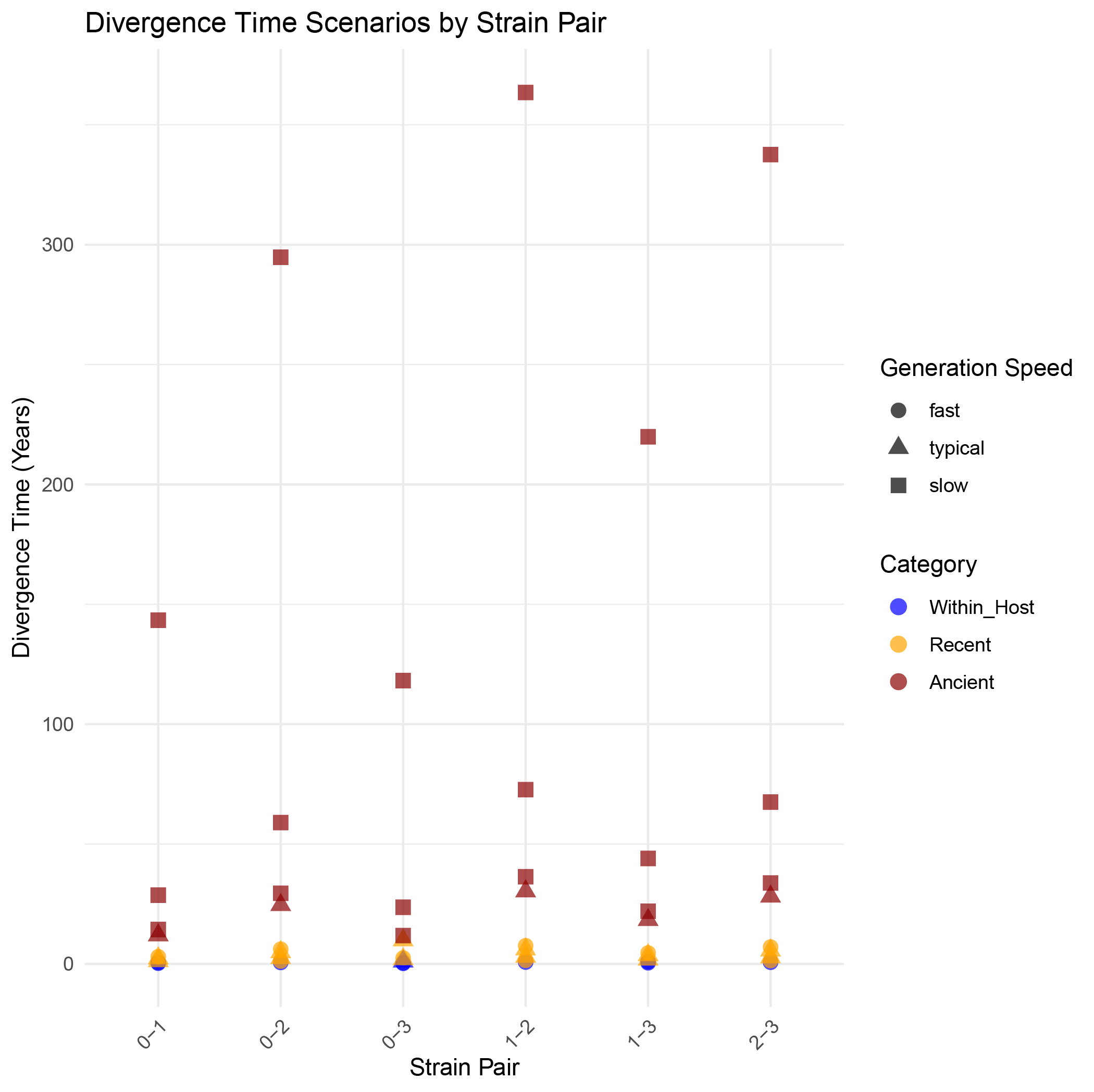

**Supplementary Figure 4. Divergence times between strains of SGB4820 under different parameter setting scenarios.** The shape of the dot represents its corresponding parameter class (fast, typecal, or slow), and the color of the dot represents its inferred evolutionary type (within_host, recent, or ancient).

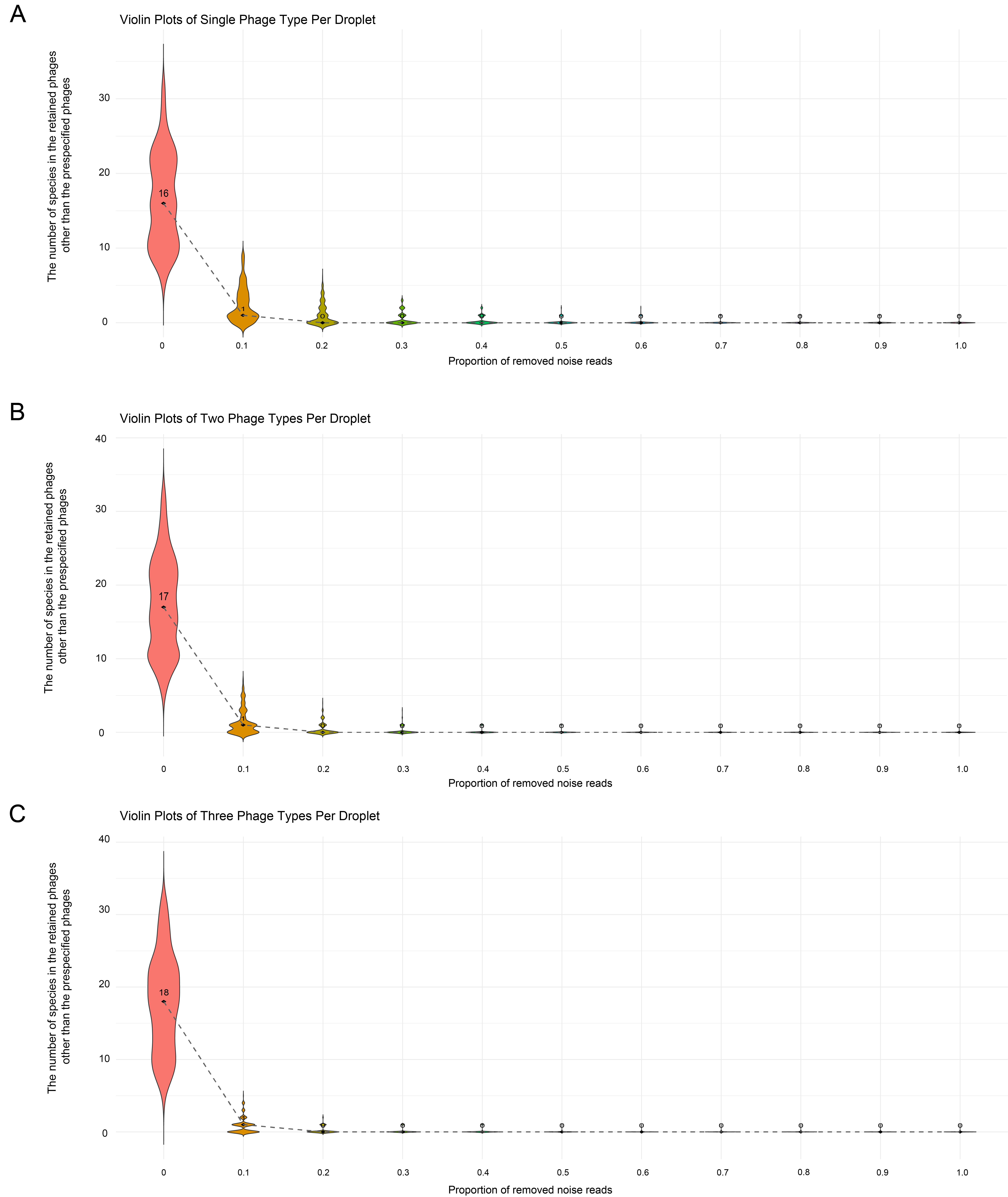

**Supplementary Figure 5. The number of non-preset phages in the simulated droplets under different phage sequence filtering thresholds**.

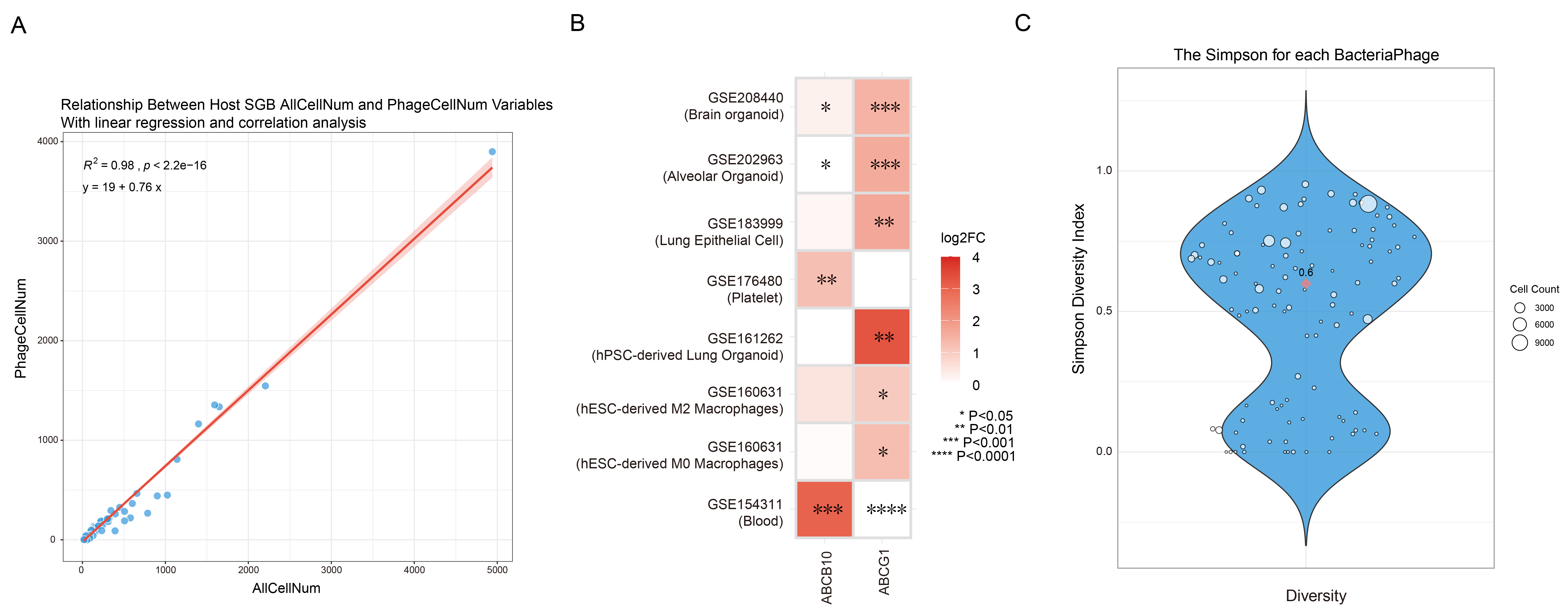

**Supplementary Figure 6.** (**A**) Correlation between total cell number and the count of cells infected by phages of all species. (**B**) Heatmap of ABCB10 and ABCG1 expression changes across SARS-CoV-2 transcriptomic datasets. Color intensity indicates log2 fold change (log2FC), with white for zero and red for higher values. (**C**) The Simpson diversity index of different phages infecting host bacteria.

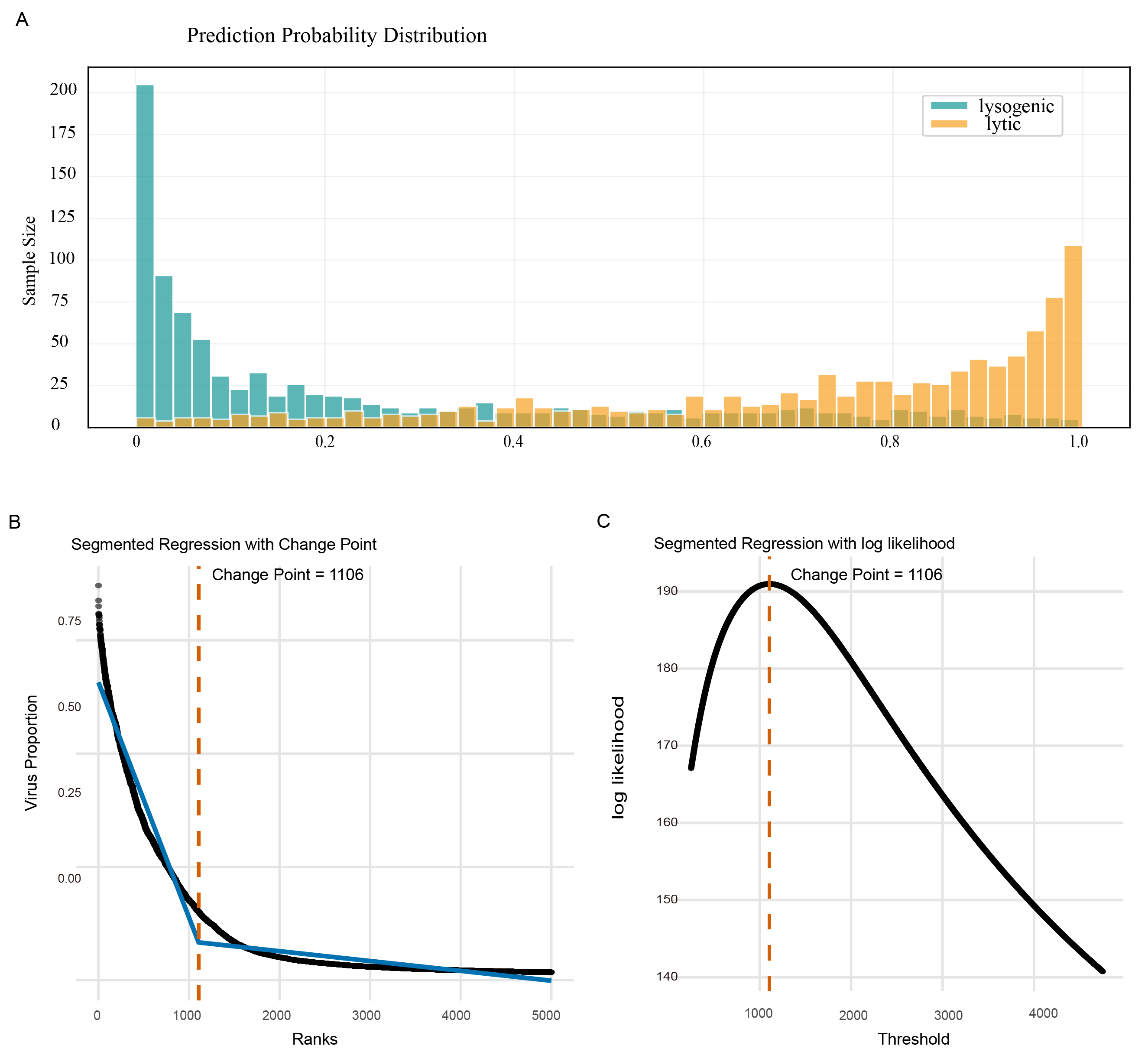

**Supplementary Figure 7.** (**A**) Density distribution of predicted lytic sequences in lysogenic phages and lytic phages. (**B**-**C**) By analysis the distribution of phage read proportions across all infected cells, a change point was found using log likelihood, and its corresponding phage reads proportion could be determined (17.15%).

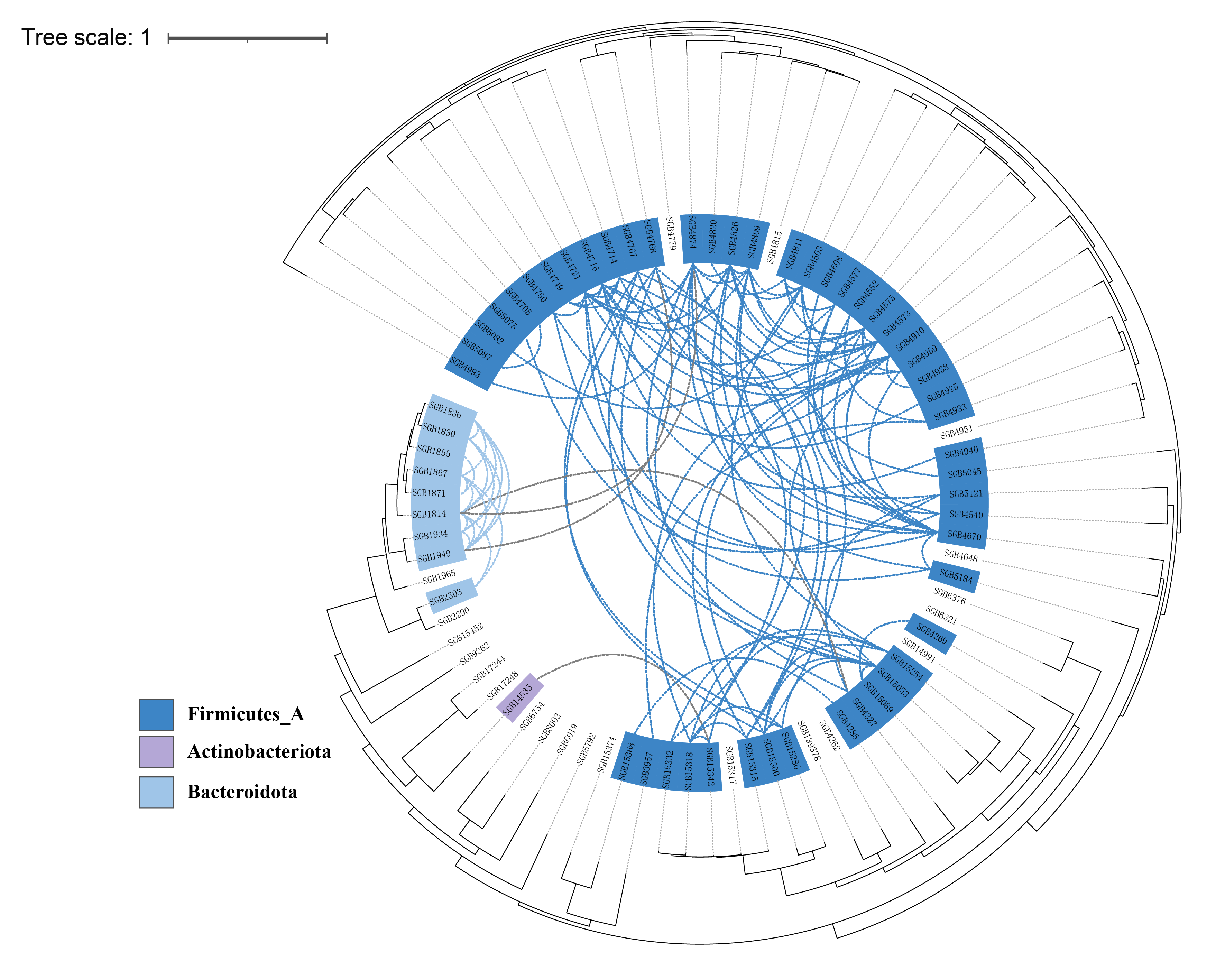

**Supplementary Figure 8. The horizontal gene transfer events among different species.**
