## Supplementary Materials and Methods for "MetaSAG: A Tool for Multi-level Exploration and Taxonomic Analysis of Microbial Single-Amplified Genomes"

**This document provides a detailed record of all the methods used in this work.**

**1. Construction of species phylogenetic trees**

We firstly converted the FASTA file of each genome into an Anvi’o contigs database (.db) using the anvi-script-FASTA-to-contigs-db program, which integrates sequence data, open reading frame predictions, k-mer frequencies, and functional/taxonomic annotations. All databases were then summarized into a central “links information file” to facilitate batch processing. Then, conserved genes were systematically identified across all 92 bacterial genomes using Anvi’o’s built-in hidden Markov model (HMM) search. The analysis detected 71 single-copy core genes in the bacterial domain, along with all major ribosomal RNA subunits (23S, 28S, 5S, 16S, 18S, 12S).

To reconstruct a robust phylogeny, six highly conserved single-copy ribosomal proteins (L1–L6) were selected as markers. Their amino acid sequences were extracted from each genome using anvi-get-sequences-for-hmm-hits, retaining only the best hit per gene to ensure strict single-copy status. The six protein sequences from each genome were concatenated in a fixed order to generate a “joint sequence” representing its core genetic signature, resulting in a matrix of 92 joint sequences.

Phylogenetic tree inference was performed with anvi-gen-phylogenomic-tree based on the joint sequence matrix, and the final tree was visualized using the iTOL web platform.

**2. Analysis of Metabolic Pathways**

This study utilized the UniProt database for functional annotation. To ensure both representativeness and sequence diversity, we selected the UniRef90 dataset as the protein reference library. UniRef90 clusters were generated using CD-HIT, with a threshold of 90% sequence identity and 80% coverage, and each cluster is represented by a single sequence.

DNA reads from each sample were aligned against a custom UniRef90 protein database using Diamond blastx, which translates reads in six frames prior to alignment, making it suitable for untranslated metagenomic reads. The alignment results were compiled into a "UniRef90 ID–cell barcode" read count matrix. To ensure data reliability, cells with fewer than five reads aligned to the UniRef90 database were removed from downstream analyses.

Based on the taxonomic classification previously assigned to each droplet (e.g., bacterial species or genus), UniRef90 alignment results from all droplets belonging to the same taxon were extracted and merged, generating taxon-specific gene family abundance profiles. These UniRef90 gene families were then mapped to MetaCyc metabolic pathways following the HUMAnN2 approach, via two reliable mapping strategies: (1) direct mapping, using UniProt accession numbers that link MetaCyc reactions to UniRef90 clusters; and (2) indirect EC-number bridging, which connects MetaCyc reactions to UniRef90 identifiers through shared Enzyme Commission numbers. A pathway was considered present in a given taxon only if its coverage—defined as the proportion of essential reactions detected—reached ≥50%, thereby minimizing false positives due to fragmented genes or random alignments.

**3. Strain classification**

This study utilized high-precision SNAP alignment to map single-cell short reads to their corresponding species reference genomes, generating BAM files. A standardized pipeline was employed for SNP identification and rigorous filtering: (1) integration of all single-cell BAM files to generate locus information; (2) variant calling based on a haploid model; (3) specific extraction of SNP variants; and (4) a triple-filtering approach using quality scores (QUAL>30), coverage (present in ≥5% of cells), and allele frequency (supported by ≥1% of cells) to obtain a high-confidence SNP matrix, followed by the removal of low-quality single-cell data.

The genotype of each cell at each SNP locus was encoded into a ternary state (1: reference, -1: variant, 0: missing/low-quality). After normalizing the encoded matrix, genetic distances between cells were computed to construct a distance matrix. Ward's hierarchical clustering was applied to generate a phylogenetic tree, and a uniform distance threshold (2.7) was used for horizontal cutting to delineate discrete strain types. Concurrently, UMAP dimensionality reduction was performed for visualization, confirming the separation of strain types.

Ultimately, clear intraspecific strain differentiation was identified across all 21 species, and the level of genetic variation for each species was quantified by calculating the "genome length/total SNP count" metric.

**4. Strain Evolutionary Analysis**

***4.1 Strain Genotype Extraction and Binary Matrix Construction***

To investigate the evolutionary relationships among bacterial strains within each SGB (Species-level Genome Bin), we developed a comprehensive analysis pipeline. First, single nucleotide polymorphism (SNP) data were extracted from the filtered SNP matrix (SGB*_SNPpd.txt) alongside strain assignment information (StrainCells.txt). For each strain, cells belonging to the same strain were identified based on the strain assignment file. Allele frequencies were calculated for each SNP position using the formula:

$$Frequency=\frac{Number of cells with alternate allele (value=1)}{Total cells with valid genotypes (values \in\{-1, 0, 1\})}$$

where -1 represents the reference allele, 0 indicates missing or ambiguous data, and 1 denotes the alternate allele. A binary genotype matrix was then constructed by applying a threshold of 0.5, where SNP positions with allele frequencies >0.5 were encoded as 1 (variant) and those ≤0.5 as 0 (reference/major allele). This conservative threshold ensured that only high-confidence variants were included in subsequent analyses.

***4.2 Phylogenetic Tree Reconstruction***

To infer evolutionary relationships among strains, phylogenetic trees were constructed using the neighbor-joining method. Pairwise genetic distances between strains were calculated using Jaccard distance, which is particularly suitable for binary genotype data:

$$D_{ij}=1-\frac{a}{a+b+c}$$

where aa represents the number of SNP positions where both strains i and j have the variant allele (1), bb represents positions where only strain i has the variant, and cc represents positions where only strain j has the variant. The distance matrix was computed using the dist() function in R with method="binary". Phylogenetic trees were reconstructed from the distance matrix using the neighbor-joining algorithm implemented in the ape package (version 5.7-1) in R. Branch lengths were optimized, and negative branch lengths (when present) were set to a minimal positive value (0.001) to ensure biological interpretability. Tree visualization was performed using the plot.phylo() function with midpoint rooting.

***4.3 Divergence Time Estimation***

Divergence times between strain pairs were estimated using a molecular clock approach. The number of SNP differences between strains was calculated as:

$${SNP}_{diff}=D_{ij}\times G$$

where *D_ij_* is the genetic distance between strains *i* and *j*, and *G* is the estimated genome size (5 Mb, typical for gut bacteria). Divergence time in years was then estimated using:

$$T=\frac{{SNP}_{diff}}{2\times\mu\times G}$$

where μμ represents the mutation rate per site per generation. We employed a mutation rate of 5×10^−7^ mutations per site per generation, based on empirical measurements for gut bacteria (Garud et al., 2019). Generation time was set to 2 hours, reflecting typical bacterial division rates in the gut environment. The final divergence time in years was converted from generations considering 4380 generations per year (2 hours/generation). To account for parameter uncertainty, we performed sensitivity analyses using three mutation rate scenarios: conservative (1×10^−5^), moderate (5×10^−6^), and liberal (1×10^−6^).

***4.4 Identification of Key SNPs***

Key SNPs that differentiate strains were identified using two complementary approaches:

***4.4.1 Strain-specific SNPs*:** For each strain, we identified SNPs that were predominantly present in that strain but absent or rare in others. A SNP was considered strain-specific if it met the following criteria: (1) Allele frequency >0.6 in the focal strain; (2) Maximum allele frequency <0.4 in all other strains

These thresholds were optimized to balance sensitivity and specificity in identifying strain-defining variants.

***4.4.2 Highly differentiated SNPs:*** To identify SNPs that contribute most to strain differentiation, we calculated the allele frequencies across all strains for each SNP position:

top 20 were designated as highly differentiated SNPs.

**5. Phage simulation data**

Given the limitation of single-cell genomic amplification techniques in capturing the complete genome within droplets, we simulated the genomic mixing scenarios between the top ten dominant bacterial species and their respective top five phages with the highest annotated abundance. Within individual droplets, we primarily explored the following two aspects: first, the accuracy (or confounding) of annotation when a single bacterium is parasitized by one or multiple phages; and second, the distribution of genomic reads when multiple phages parasitize a single bacterium.

We assumed that each droplet contained only the genome of one bacterial species, with the number of parasitic phage species ranging from 1 to 3. For each parasitic scenario, 300 single-amplified genome droplets were simulated, with each phage species present in 1, 10, or 20 copies per droplet. Paired-end short reads were simulated for each amplified droplet and subsequently annotated. Some non-preset phages were detected in the annotation results of individual droplets, which could be attributed to genomic sequence homology among phages or the presence of phage fragments within the bacterial genome itself. Phage sequences were identified based on annotations from the mpa_vJun23_CHOCOPhlAnSGB_202403_VSG.fna.

To eliminate these phages that caused random annotation confusion, we applied a proportional threshold based on the cumulative distribution of phage reads, removing phages with relatively low read counts. The results revealed that removing phages whose reads accounted for the top 20% of the total phage reads within a droplet effectively eliminated all non-preset phage labels. This proportional threshold remained unaffected by the number of phage species or their copies within the droplet. Additionally, we required that the retained phages in the droplet have read counts ≥80 to ensure the reliability of subsequent analyses.

**6. Simpson Diversity Index Analysis for Viral Host Range Quantification**

To quantitatively assess the host range of bacteriophages based on single‑cell sequencing data, we adapted the Simpson diversity index—a metric commonly used in microbial ecology to evaluate species richness and evenness in community analyses. The index was employed here to characterize the breadth and specificity of phage–host interactions.

The Simpson index (D) is calculated as:

D = 1 − Σ *p*ᵢ²

where *p*ᵢ represents the proportion of infections attributed to the i^th^ bacterial host. The value of D ranges from 0 to 1. A value close to 0 indicates that the phage infects only a narrow set of hosts (high host specificity), whereas a value approaching 1 reflects a broader and more even distribution of infections across multiple host taxa. Compared to simple host‑species counting, the Simpson index incorporates both the richness of host types and the relative infection intensity across hosts.

To identify the core phage–host interactions that dominate the observed diversity, we further calculated the cumulative contribution of hosts to the Simpson index for each phage. Specifically, for a given phage, all its hosts were ranked in descending order of infection proportion (*p*ᵢ). The cumulative contribution of the top k hosts to the index was computed sequentially until reaching 90% of the total Simpson index value. Hosts included up to this 90% threshold were regarded as the major contributors to phage host diversity.

Only phage–host pairs meeting the following criteria were retained for downstream analysis: (1) The host belongs to the set of core contributors (cumulative contribution ≥90% of the Simpson index); (2) The number of infected cells for that phage–host pair is ≥10.

**7. Development and Training of MetaK-Lytic: A Few-Shot Learning for Phage Lysis Ability Prediction**

To address the challenge of classifying phage lifestyles (lytic vs. non-lytic) under limited labeled data, we designed and implemented a novel meta-learning framework based on a Memory-Augmented Neural Network (MANN). The core innovation of this framework is the incorporation of an external memory module, which enables the model to rapidly assimilate information from few samples and generalize to new categories, thereby overcoming the tendency of traditional deep learning models to overfit and their inability to adapt dynamically in small-sample scenarios.

The data was downloaded using the following code:

\# Download all complete phage genomes with "phage" in their names from RefSeq (FASTA format)

ncbi-genome-download \

--formats fasta \ # Output format: fasta/genbank/assembly-report, etc. "all" can be specified

--assembly-levels complete \ # Download only fully assembled genomes

--genera phage \	# Match genus names containing "phage"

--fuzzy-genus \ # Fuzzy matching (critical: otherwise only genus names exactly equal to "phage" are matched)

--parallel 16 \ # Parallel downloads (speed up, adjust the number based on network speed)

viral	# Core taxonomic group: phages belong to the viral category

#### *7.1 Meta-Learning Task Formalization: Episodic Training*

We adopted an "episodic training" paradigm, organizing the training process into a series of episodes. Each episode simulates a few-shot learning task. Formally, an episode EE is defined as a small dataset
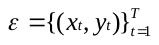
, where:

*x_t_* is the k-mer feature vector of a phage sample (loaded from the pre-processed .npy files) at timestep tt.

*y_t_*∈{0,1} is the corresponding phage lifestyle label (0 for lytic, 1 for temperate).

*T* is the number of samples in the episode.

A critical design to force the model to rely on the external memory rather than its static weights is the use of **temporally-shifted input** and **random label remapping**.

**Temporally-shifted input**: The input at timestep *t* is the tuple (*x_t_*, *y_t_*_−1_). At the first timestep (*t*=1), where no previous label exists, the input is (*x*_1_, 0). The model is tasked at each step *t* to predict the current label *y_t_*.

**Random Label Remapping**: The mapping between the biological category and the numerical label (0/1) is randomly permuted at the start of each new episode. For instance, 'temperate' might be labeled 1 in one episode and 0 in the next.

This setup prevents the model from slowly encoding fixed label representations into its weights. Instead, it must learn a universal strategy: to dynamically bind the currently observed sequence features *x_t_* with the label *y_t_* (provided in the next input) and store this association in the external memory for subsequent retrieval and prediction.

#### *7.2 Model Architecture: MANN with LRUA Access Mechanism*

Our model is based on the Neural Turing Machine architecture, enhanced with a **Least Recently Used Access (LRUA)** memory writer, which is a pure content-based addressing mechanism well-suited for rapid binding of new information.

***7.2.1 Controller***

The controller, implemented as a Long Short-Term Memory (LSTM) network due to its ability to handle sequential dependencies, serves as the main processing unit. It takes the input (*x_t_*, *y_t_*_−1_) (with *y_t_*_−1_ passed through an embedding layer) and updates its hidden state. The controller's output vector is used to generate a key vector *k_t_*, which governs subsequent memory operations.

***7.2.2 External Memory Matrix***

The external memory is represented as a matrix *M_t_* of size *N*×*W*, where *N* is the number of memory locations and *W* is the vector dimension at each location. This matrix acts as a dynamically addressable workspace.

***7.2.3 Reading Mechanism***

To read from memory, a read weight vector
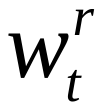
 is computed based on the cosine similarity between the key vector *k_t_* and each row *M_t_*(*i*) in the memory matrix:

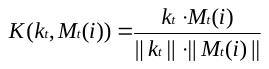

The read weights are obtained by applying a softmax over these similarities:

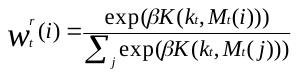

The retrieved memory vector *r_t_* is then the weighted sum:

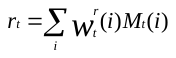

This vector *r_t_* is concatenated with the controller's output and passed to a softmax classification layer for predicting *y_t_*. It is also fed back into the controller as input for the next timestep.

***7.2.4 Writing Mechanism (LRUA)***

The LRUA mechanism determines where to write new information (the key vector *k_t_*) by prioritizing either the most recently used (MRU) or least recently used (LRU) memory locations.

The model maintains a usage weight vector
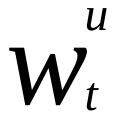
, updated at each timestep to track how frequently each location has been used.

The least-used weight vector
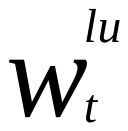
 is derived from
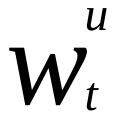
, setting the values for the *m* least-used locations to 1 and others to 0.

The write weight vector
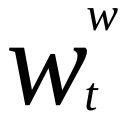
 is a convex combination of the previous read weights
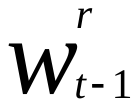
 and the previous least-used weights
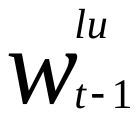
, controlled by a learnable gating parameter *α*:

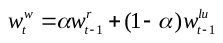

This allows the model to choose between updating a recently read location (potentially reinforcing or correcting information) or writing to a rarely used, potentially empty location (protecting recently acquired knowledge).

Before writing, the memory locations identified by
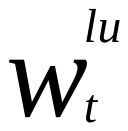
 are cleared. The new information is then added to the memory:

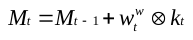

#### *7.3 Meta-Learning Training Procedure*

The MANN was trained over a large number of episodes. In each episode, a small subset of phage classes (e.g., 5 classes, simulated via label remapping to create a multi-task environment even for binary classification) was randomly sampled from the training set.
The loss (categorical cross-entropy) was calculated at each timestep for the prediction
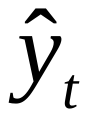
 against the true label *y_t_*. The total loss for an episode was the average of these step-wise losses.
The objective was not merely to minimize loss on a fixed dataset but to minimize the expected loss across the distribution of possible episodes
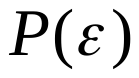
:

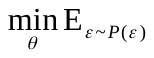
[ℒ(ε; θ)]

where *θ* represents the model parameters. This encourages the model to learn a general-purpose "binding-and-retrieval" strategy applicable across tasks.

The external memory was reset at the end of each training and testing episode to prevent interference from unrelated information across different tasks, ensuring the model started each new task with a clean working memory.
